# All-trans retinoic acid induces durable tumor immunity in IDH-mutant gliomas by rescuing transcriptional repression of the CRBP1-retinoic acid axis

**DOI:** 10.1101/2024.04.09.588752

**Authors:** Aparna Rao, Xiaoran Zhang, Anthony R. Cillo, Jonathan H. Sussman, Poorva Sandlesh, Antonio Corral Tarbay, Arka N. Mallela, Carly Cardello, Katharine Krueger, Jessica Xu, Alex Li, Jason Xu, Jonathan Patterson, Ebrar Akca, Angelo Angione, Emade Jaman, Wi Jin Kim, Jordan Allen, Abhishek Venketeswaran, Pascal O. Zinn, Robert Parise, Jan Beumer, Anette Duensing, Eric C. Holland, Robert Ferris, Stephen J. Bagley, Tullia C. Bruno, Dario A.A. Vignali, Sameer Agnihotri, Nduka M. Amankulor

**Affiliations:** Department of Neurological Surgery, University of Pittsburgh, Pittsburgh, PA; Hillman Cancer Center, University of Pittsburgh Medical Center, Pittsburgh, PA; Fred Hutchinson Cancer Center, Seattle, WA; Department of Neurological Surgery, University of California Los Angeles, Los Angeles, CA; University of Pittsburgh School of Medicine, Pittsburgh, PA; Rangos Research Center, UPMC Children’s Hospital of Pittsburgh, Pittsburgh, PA; Department of Pharmaceutical Sciences, University of Pittsburgh, Pittsburgh, PA; Department of Otolarnyngology, Head & Neck Division, University of Pittsburgh, Pittsburgh, PA; Department of Neurological Surgery, University of Washington, Seattle, WA; Department of Immunology, University of Pittsburgh, Pittsburgh, PA; Tumor Microenvironment Center, UPMC Hillman Cancer Center, Pittsburgh, PA; Division of Hematology Oncology, Department of Medicine, University of Pennsylvania, Philadelphia, PA; Department of Neurological Surgery, University of Pennsylvania, Philadelphia, PA; Genomics and Computational Biology Graduate Group, University of Pennsylvania, Philadelphia, PA

## Abstract

Diffuse gliomas are epigenetically dysregulated, immunologically cold, and fatal tumors characterized by mutations in isocitrate dehydrogenase (IDH). Although IDH mutations yield a uniquely immunosuppressive tumor microenvironment, the regulatory mechanisms that drive the immune landscape of IDH mutant (IDHm) gliomas remain unknown. Here, we reveal that transcriptional repression of retinoic acid (RA) pathway signaling impairs both innate and adaptive immune surveillance in IDHm glioma through epigenetic silencing of retinol binding protein 1 (RBP1) and induces a profound anti-inflammatory landscape marked by loss of inflammatory cell states and infiltration of suppressive myeloid phenotypes. Restorative retinoic acid therapy in murine glioma models promotes clonal CD4^+^ T cell expansion and induces tumor regression in IDHm, but not IDH wildtype (IDHwt), gliomas. Our findings provide a mechanistic rationale for RA immunotherapy in IDHm glioma and is the basis for an ongoing investigator-initiated, single-center clinical trial investigating all-trans retinoic acid (ATRA) in recurrent IDHm human subjects.

## INTRODUCTION

Mutations in isocitrate dehydrogenase 1 and 2 (*IDH1* and *IDH2*) are frequent genetic drivers of human cancers^1^. These mutations are characteristic hallmarks of a subset of primary brain glial malignancies including diffuse low-grade gliomas (DLGGs) and secondary glioblastomas (sGBM)^2^. Such mutations are found in ∼80-90% of DLGGs and nearly all sGBMs^3,4^. Wild-type isocitrate dehydrogenase catalyzes the oxidative carboxylation of isocitrate to alpha-ketoglutarate (α-KG), whereas mutant IDH (mIDH) enzymes, most commonly featuring an arginine to histidine substitution at codon 132 (R132H), further catalyze the conversion of α-KG into the D-2-hydroxyglutarate (D-2HG) oncometabolite^5^. IDH mutation-driven oncogenesis is marked by epigenetic reprogramming through hypermethylation of genomic CpG and histone loci and is a direct consequence of D-2HG-mediated inhibition of α-KG-dependent hydroxylases and demethylases (including JmjC histone demethylases)^6,7^. The consequence of epigenetic reprogramming of IDH-mutant (IDHm) gliomas is a distinct methylomic signature known as the glioma CpG Island Methylator Phenotype (g-CIMP)^8^. The net oncogenic properties of IDHm gliomas can be summarized by its transcriptional output.

IDHm and IDH wild-type (IDHwt) gliomas are oncological and genetically disparate entities, and these differences are captured by salient, sometimes pathognomonic, histologic, radiographic, and clinical characteristics. For instance, IDHm gliomas may exist in prolonged dormancy prior to high-grade transformation, whereas IDHwt gliomas are usually progressive and highly invasive at diagnosis^9–11^. The dormant phase seen in IDHm glioma has raised the possibility that prolonged impairment of tumor immune surveillance and deficient innate immune function is a required for high-grade transformation. Indeed, previous studies have identified unique loss of effector tumor infiltrating lymphocytes^12–14^, increased infiltration of suppressive myeloid cell populations, and global loss of inflammatory signaling mediated by STAT1/IFN-γ, and diminished cytolytic activity of natural killer (NK) cells in IDHm tumors^15–17^. Epigenomic imprinting in IDHm glioma clearly regulate cell-intrinsic expression of anti-tumor immune checkpoints, in part by methylation of activating NKG2D ligands whose stress-induced expression target cells for lysis by innate lymphoid cells; however, no overarching regulatory immune suppression has been identified^16^.

Several investigators have hypothesized that tumor cell-intrinsic and extrinsic immune dysfunction in IDHm gliomas are mediated by specific non-stochastic epigenetic events downstream of 2-HG production^18,19,16,20,21^. Using multimodal transcriptional analysis of IDHm glioma we show that marked transcriptional repression of *RBP1*, a chaperone gene required for retinoic acid (RA) synthesis, is a master regulator of innate immune dysfunction in IDHm glioma. Importantly, widespread immune dysfunction in IDHm gliomas can be reversed with exogeneous repletion of RA using ATRA. Remarkably, ATRA repletion therapy remodels the tumor immune microenvironment (TIME) to permit anti-tumor immunity. Importantly, we show that ATRA enhances immunogenicity of IDHm gliomas through cell-autonomous induction of activating NK cell ligands and clonal expansion of novel CD4^+^ TCR repertoire. These findings establish a paradigm that unifies previously disparate elements of immune suppression in IDHm glioma (i.e., NK ligand repression, loss of lymphoid cells, and infiltration of suppressive myeloid ontogenies).

## RESULTS

### Loss of RA signaling is a hallmark of myeloid dysregulation in IDHm glioma

Arrested myeloid differentiation is characteristic of the tumor microenvironment (TME) of all gliomas, but is particularly prominent in IDHm gliomas^22^. To identify genotype-specific pathways that mediate myeloid homeostasis in IDHm glioma, we characterized gene expression differences between IDHm and IDHwt gliomas from The Cancer Genome Atlas (TCGA) low-grade glioma (LGG) patient cohort and extracted significantly altered immune regulatory genes **(Figures 1a-b)**. Our analysis demonstrated stark genotype-dependent differences, including 393 differentially-expressed genes (DEGs) matching genes within the *Leukocyte Differentiation* Gene Ontology gene set^23^ (**Figure 1a**). *RBP1,* a retinoid pathway gene whose protein product shuttles the vitamin A derivative, retinol, to retinol dehydrogenases (RDHs) during ATRA biosynthesis, emerged as one of the most downregulated leukocyte differentiation-related genes in IDHm gliomas (**Figure 1b**). Next, we explored the relationship between epigenetic dysregulation and gene expression of the *RBP1* locus by characterizing the extent of CpG island hypermethylation of *RBP1* and identified a statistically significant inverse correlation between *RBP1* methylation and transcription (LGG TCGA database) (**Figure 1c**). *RBP1* transcriptional repression was present in both 1p/19q co-deleted and non-co-deleted patient samples (**Figure 1d**), inferring a dependence on the IDH genotype rather than cell-of-origin. *RBP1’s* protein product, CRBP1, was nearly undetectable in Western blots of IDHm patient-derived glioma stem cells (GSCs) compared to IDHwt GSCs (**Figure 1e**), consistent with prior reports^24^. Most importantly, mass spectroscopic analysis of ATRA revealed a stunning reduction (>800-fold) in intracellular ATRA in IDHm glioma cells and directly linked transcriptional repression of *RBP1* with abrogation of ATRA synthesis (**Figure 1f**).

**Figure 1.**
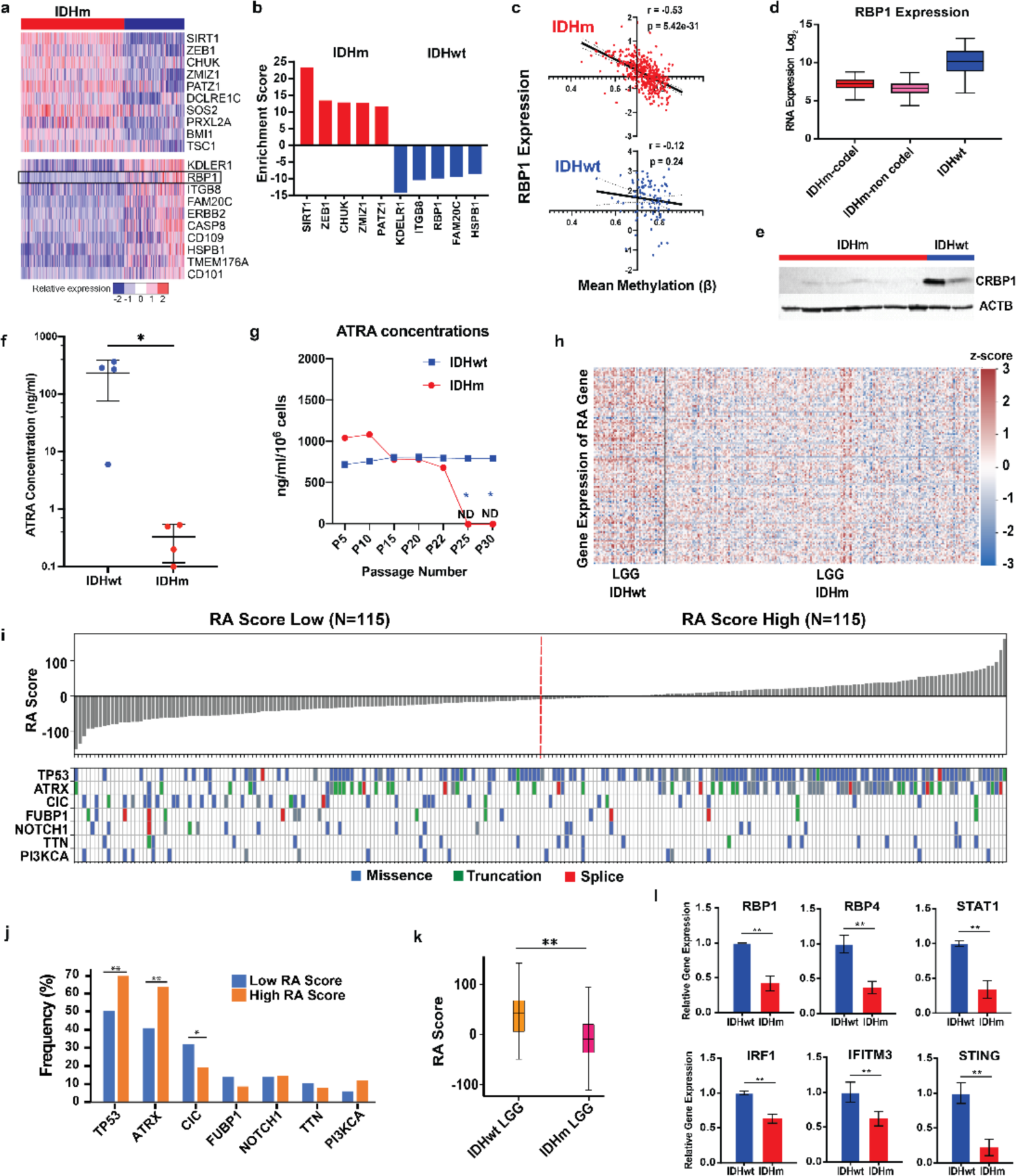
Loss of RA signaling is a hallmark of myeloid dysregulation in IDHm gliomas. (a) Heatmap of gene expression from RNA-seq of selected genes from the *Leukocyte Differentiation* Gene Ontology gene set across patients from the TCGA-LGG cohort. Gene expression is normalized by row. (b) Quantitative representation of top differentially expressed genes based on enrichment score. (c) Correlation plot of *RBP1* methylation and gene expression in IDHwt and IDHm samples from the TCGA-LGG database, with Pearson’s correlation coefficient (*r)* shown. (d) *RBP1* expression level in both 1p19q co-deleted and non-co-deleted IDHm patient samples from the TCGA-LGG cohort. (e) CRBP1 protein expression level as assessed by Western blot in 7 IDHm samples and 2 IDHwt samples from patient-derived tumors. (f) Mass spectrometry quantification of intracellular ATRA concentration (ng/ml) in lysates of GSC lines of 3 IDHm and 4 IDHwt tumors. * p < 0.05 calculated by two-tailed Student’s *t*-test. (g) Mass spectrometry quantification of intracellular ATRA concentration (ng/ml) in lysates of 3 IDHm 3 IDHwt astrocyte cell lines at different passages. * p < 0.05 calculated by two-tailed Student’s *t*-test. (h) Interrogation of the TCGA-LGG dataset using a published 132-gene signature derived from differentially expressed genes in RA-treated human monocytes. (i) Schematic depicting the association between high and low RA score and types of secondary mutations at multiple genes of interest. (*n* = 115 patients defined as having a high and low RA score each based on median RA myeloid score). (j) Bar plot of associations between secondary mutations and RA score category. Values indicate frequency of specified secondary mutation within the low RA score and high RA score categories respectively. * p < 0.05, ** p < 0.01 via two-tailed *z*-score test between two proportions. (k) Average RA scores of IDHwt and IDHm LGG samples in within the TCGA-LGG cohort. ** p < 0.01, calculated by two-tailed Student’s *t* test. (l) Expression levels of RA inducible genes between ATRA-treated IDHm and IDHwt astrocyte cell lines (*n* = 3) assessed by qPCR. ** p < 0.01 via two-tailed Student’s *t*-test.

Epigenetic reprogramming is dependent on progressive cell division and correlates with passage of cells *in vitro*^25–27^. To establish whether loss of ATRA is a feature of epigenetic consolidation, we passaged well-characterized astrocyte cell lines transfected with a lentiviral vector encoding wildtype IDH1 (IDHwt) or IDH1-R132H (IDHm)^8^. We then measured intracellular ATRA concentration using mass spectrometry after every fifth passage between p0 and p30. IDHwt astrocytes maintained stable ATRA concentration over all 30 passages (∼600ng/ml/10^6^ cells), whereas IDHm astrocytes demonstrated a dramatic decline in ATRA concentrations beginning at p10 and progressing to undetectable concentration levels by p25 (**Figure 1g**). Thus, loss of ATRA synthesis is a direct consequence of oncogenic transcriptional repression following induction of epigenetic memory.

RA is associated with differentiation of immature/suppressive myeloid cell states into mature phagocytic macrophage ontologies. To this end, we quantified the relative presence or absence of ATRA-specific myeloid transcriptional signatures by leveraging a previously validated 132-gene signature derived from differentially expressed genes in RA-treated human monocytes^28^. Using the sum of TCGA-derived normalized z-scores across the 132 RA myeloid genes as a composite myeloid RA score for each LGG sample, we noted a significantly lower myeloid RA score in IDHm gliomas compared to IDHwt (**Figure 1h-k**). Interestingly, myeloid-related ATRA scores were associated with other recurrent genomic alterations. *TP53* and *ATRX* mutations were more likely to occur in the high RA score category, while mutations in *CIC* were more likely to occur in the low RA score category (**Figure 1i-j**). Although IDH mutation is the primary regulator of RA dysfunction, these data suggest that secondary mutations can modulate the degree of RA-mediated myeloid dysfunction within IDHm tumors. Lastly, interrogation of RA-induced pro-inflammatory genes via quantitative PCR (qPCR) in IDHm and IDHwt astrocytes also revealed lower expression of known RA-induced genes and revealed downregulation of interferon-related (*IFIT1, STAT1, STING*) genes that alter myeloid function in IDHm astrocytes. Taken together, these data provide direct evidence of transcriptional repression of RA signaling in IDHm gliomas and suggest that myeloid cell fates might be regulated by RA in human glioma.

### ATRA induces distinct leukocyte-activating transcriptional signatures in IDHm astrocytes

To establish a more thorough understanding of ATRA-induced gene expression and to decipher biological pathways activated in IDHm glioma following ATRA treatment, bulk RNA sequencing (RNA-seq) was performed in human IDHm and IDHwt astrocyte cell lines treated with ATRA or DMSO. We utilized well-characterized immortalized astrocyte cell lines rather than IDHm glioma cells for this purpose to minimize the effect of confounding genomic alterations^8^. We confirmed mutant IDH1-R123H status via western blot (**Extended Data Figure 1a**). Additionally, cell morphology and cellular differentiation markers were also evaluated after treatment with ATRA, DMSO, and 10% FBS (**Extended Data Figure 1b-e**). ATRA-treated cells displayed profound transcriptional changes compared to DMSO-treated counterparts in both IDHm and IDHwt cell lines, representing the primary axis of variation (**Figure 2, Extended Data Figure 2a**). Many differentially expressed genes between IDHm ATRA and DMSO-treated cells were involved in immune processes, including *ICAM1, APOE, SOCS3, CSF1,* and *IL16* **(Figure 2b**). To examine the global effect of ATRA on IDHm cell lines, we explored differentially expressed immunobiological pathways using Gene Set Enrichment Analysis (GSEA). GSEA of both ATRA- and DMSO-treated IDHm cells compared to IDHwt cells confirmed baseline repression of retinoic acid and immune-related pathways (**Extended Data Figure 2b-d**). Importantly, we observed differential responses to ATRA treatment in IDHm cells compared to IDHwt cells (**Figure 2c**). Using a linear model to identify genes differentially regulated by ATRA in IDHm cells, we discovered multiple immune-related pathways that were significantly altered (all enriched) as a consequence of ATRA treatment in IDHm conditions (**Figure 2d**). The enriched gene sets primarily include pathways representing immune cell activation and differentiation (e.g., *Myeloid Leukocyte Activation*) and immune targeting (e.g., *Leukocyte-Mediated Cytotoxicity*) (**Figure 2d-e**). Lastly, ATRA induced alterations in pathways beyond immune activation, including broad changes in metabolic activity and cell signaling in IDHm cells (**Extended Data Figure 2e**). Taken together, these results demonstrate that ATRA significantly alters the transcriptional landscape in astrocyte cell lines in an IDHm-dependent manner, specifically upregulating immune regulatory pathways and inducing leukocyte activation under IDHm conditions.

**Figure 2.**
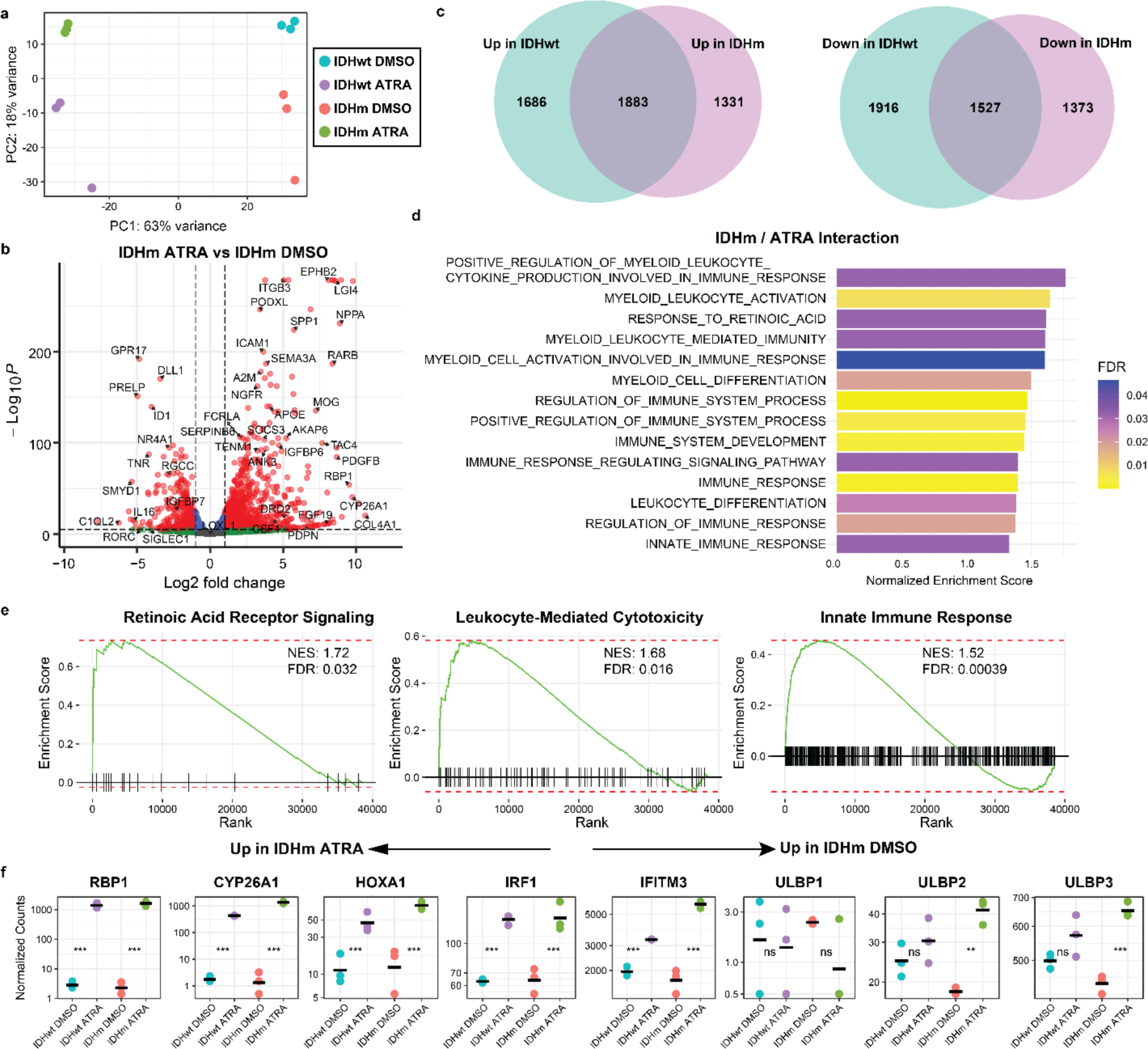
ATRA induces leukocyte-activating transcriptional signatures in IDHm astrocytes. (a) Principal component analysis of bulk RNA-seq transcriptomic profiles of IDHwt and IDHm astrocyte cell lines treated with DMSO or 1µM ATRA for 48 hours, using all genes (*n* = 3 per condition). (b) Volcano plot highlighting differentially expressed genes between ATRA-treated IDHm cell lines versus DMSO-treated IDHm cell lines. (c) Number of differentially expressed genes (DEGs) shared between IDHwt and IDHm astrocyte cell lines after ATRA treatment. DEGs defined as p-adjusted < 0.05. (d) Gene set enrichment analysis (GSEA) of Gene Ontology pathways related to retinoic acid signaling and immune response based on genes differentially expressed by interaction of IDHm and ATRA conditions. A linear model was used to rank genes by their expression change as a result of ATRA treatment specifically in the IDHm genotype beyond the additive effect of genotype plus treatment. Pathway results demonstrate upregulation of retinoic acid receptor signaling and immune responses upon ATRA treatment preferentially in IDHm astrocyte cell lines. All significant pathways (FDR <0.05) are shown. (NES, Normalized Enrichment Score; FDR, False Discovery Rate). (e) Normalized counts of selected retinoic acid response genes and NKG2D ligands from bulk RNA-seq. Statistical significance for IDHm and IDHwt cell lines between ATRA-treated and DMSO conditions is calculated via DESeq2. *** p-adj <0.001, ** p-adj <0.01, ns; not significant.

### ATRA promotes tumor cytotoxicity by activating the NKG2D immune checkpoint in an *RBP1*-dependent manner

Our group and others have reported IDH glioma-specific functional inhibition of immune-mediated cytotoxicity induced by transcriptional repression of activating NKG2D-family of NK cell ligands (NKG2DLs), including the UL-16-related *ULBP1-3*^16,29,30^. Since ATRA is a potent transcriptional activator of NKG2DLs^31^, we evaluated the extent to which ATRA induces NK-mediated cytotoxicity and the role of NKG2DLs following exposure of IDHm or IDHwt cells to ATRA using the astrocyte cell line described above. Indeed, through our RNA-seq analysis, we found that NKG2DLs *ULBP2* and *ULBP3* were exclusively upregulated upon ATRA treatment in glioma cell lines (**Figure 2f**). Upregulation of NKG2DLs was associated with markedly increased IDHm-specific NK-mediated cytotoxicity (**Figures 3a-c**). Moreover, NKG2DL upregulation exhibited distinct genotype-restriction, characterized by ATRA-induced upregulation of *ULBP1, ULBP2,* and *ULBP3* along with a robust increase in NK-mediated cytotoxicity in IDHm, but not IDHwt genotypes (**Figures 3d-f**). Importantly, ATRA-induced NK-mediated cytotoxicity was found to occur in a dose dependent manner (**Extended Data Figure 3b-c**). Moreover, the 2HG small molecular inhibitor (AG-5198) produced similar upregulation of NKG2DLs as ATRA, suggesting that transcriptional activation of NKG2DLs by ATRA is dependent on the presence of 2-HG (**Extended Data Figure 3d-e**).

**Figure 3.**
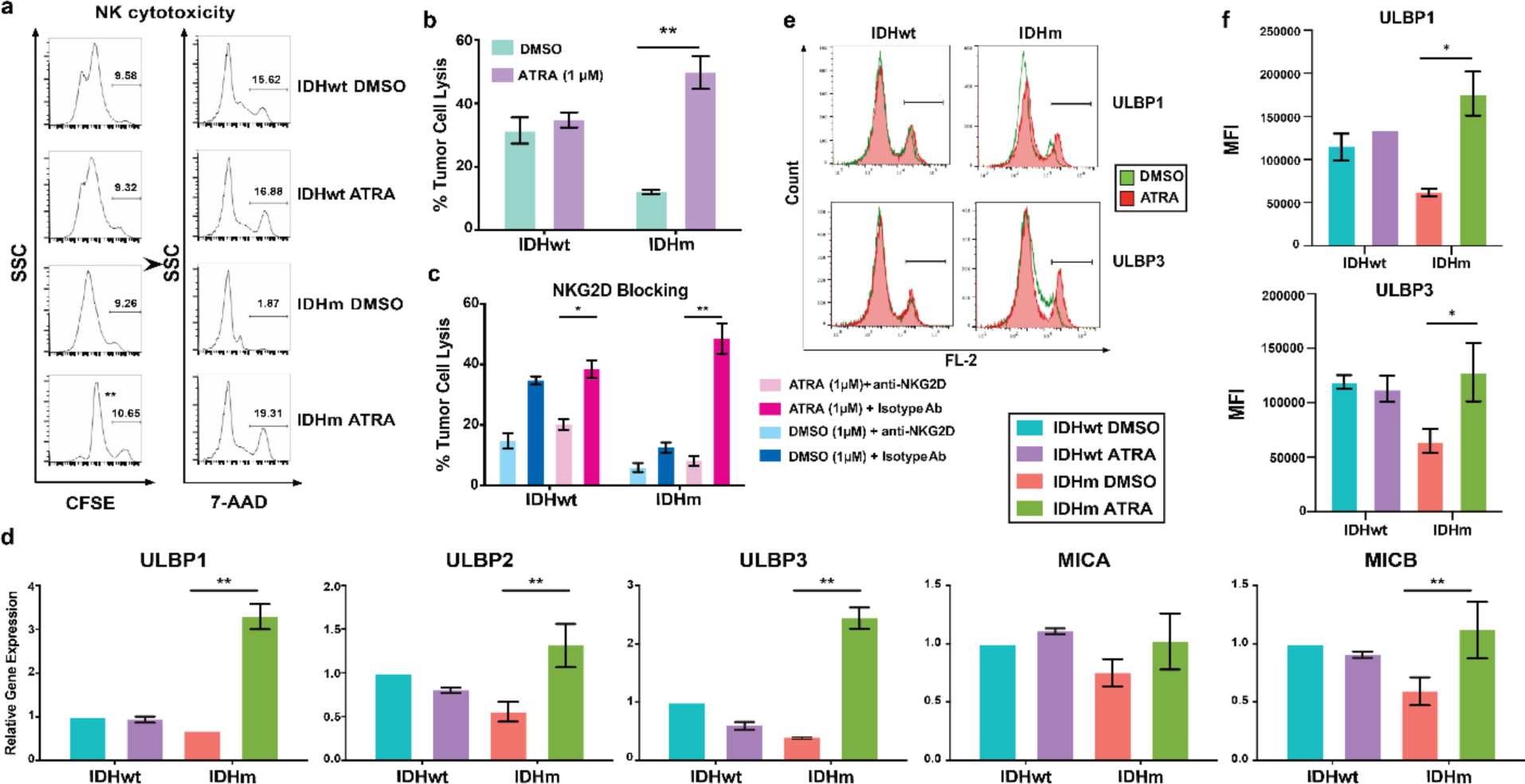
ATRA mediates NK cytotoxicity in IDHm but not IDHwt GSCs. (a) Representative flow cytometry gating strategy for assessing NK-mediated cytotoxicity across IDHwt and IDHm glioma stem cells (GSCs) upon ATRA treatment. (b) NK-mediated cytotoxicity on IDHm and IDHwt GSCs after 1 µM of ATRA or DMSO for 48 hours (*n* =3). (c) Anti-NKG2D antibody treatment reveals NKG2D dependence of ATRA-induced NK cytotoxicity. (d) Gene expression of NKG2D ligands after treatment with 1 µM of ATRA or DMSO 48 hours assessed by qPCR assay (*n* = 3). (e) Representative gating strategy for assessment of surface ULBP1 and ULBP3 expression using flow cytometry. (f) Surface expression of NKG2D ligands in GSCs after treatment with 1 µM of ATRA or DMSO for 48 hours. (b-d, f) Significance calculated via two-tailed Student’s t test. * p < 0.05, ** p < 0.01.

IDH mutations induce epigenetic repression of *RBP1* transcription which, in turn, diminishes ATRA synthesis exclusively in IDHm glioma (**Figure 1**). We sought to extend the hypothesis that retinoid production regulates immunogenicity in IDHm glioma by determining whether *RBP1* is necessary and sufficient to induce the immunomodulatory phenomena observed in IDHm gliomas. To this end, we first knocked down *RBP1* using short hairpin RNA (shRNA) in IDHwt cells. Following stable transduction and confirmation of *RBP1* knockdown (**Figure 4a**), we assessed representative hallmarks of cell-autonomous immune deficiency, including NKG2DL gene expression, NK-mediated cytotoxicity, and expression of type 1 interferon/RA-dependent gene expression. *RBP1*-silenced IDHwt cells displayed markedly lower NKG2DL expression (*ULBP1-3*) and were more resistant to NK-92-mediated cytotoxicity than their mock-transduced counterparts (**Figure 4b-d**). NK-mediated cytotoxicity in *RBP1*-silenced IDHwt cells was virtually identical to baseline IDHm conditions (**Figure 4b**), suggesting that the RA pathway maintains homeostatic thresholds of stress-induced NK cytotoxicity. Interesting, *RBP1* knockdown induced downregulation of multiple interferon and retinoid-related genes, recapitulating expression patterns of IDHm cells (**Figure 4d**).

**Figure 4.**
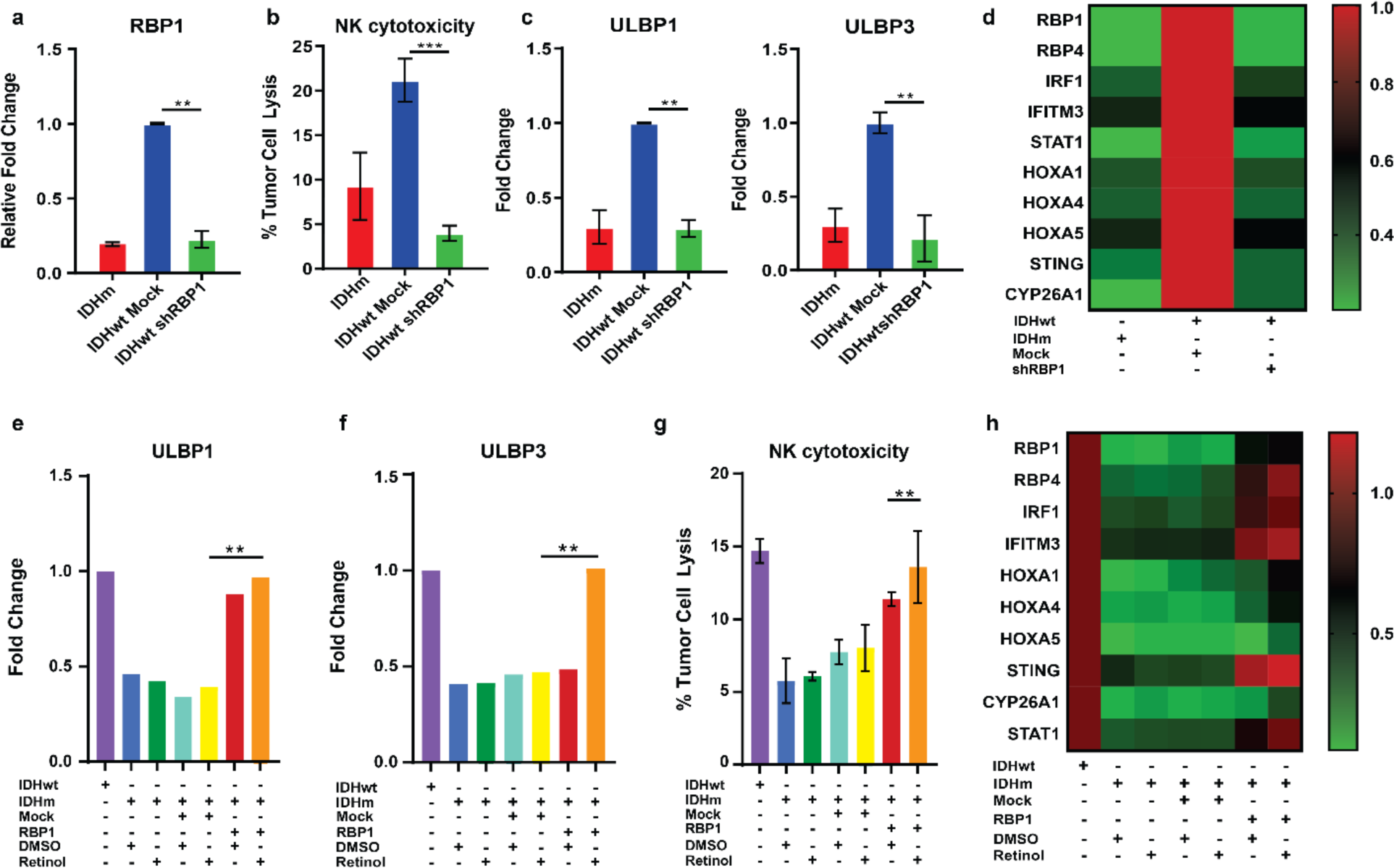
Transcriptional repression of *RBP1* is necessary and sufficient for NKG2D-dependent NK cell resistance. (a) *RBP1* gene expression in IDHm, IDHwt mock transfected, and sh*RBP1* transfected GSCs assessed by qPCR. Expression was normalized against mock-transfected IDHwt cells. (b) Flow cytometry analysis of percent of 7-AAD^+^ as a surrogated for NK-mediated cytotoxicity in *RBP1-*silenced GSCs and controls. NK:tumor cell E:T ratio of 1:10 was used. (c) Gene expression for NKG2D ligands in *RBP1**-***silenced GSCs and controls, assessed by qPCR. (d) Heatmap of relative gene expression levels corresponding to RA-dependent canonical genes and NKG2D ligands. (e-f) Gene expression for *ULBP1* (e) and *ULBP3* (f) in *RBP1**-***expressing GSCs and controls after 48 hours of treatment with 10 µM of ATRA or DMSO, assessed by qPCR. (g) Flow cytometry analysis of percent of 7-AAD^+^ as a surrogated for NK-mediated cytotoxicity in *RBP1* silenced GSCs and controls after 48 hours of treatment with 10 µM of ATRA or DMSO. NK:tumor cell E:T ratio of 1:10 was used. (h) Heatmap of gene expression levels in *RBP1-*silenced GSCs and controls corresponding to RA-dependent canonical genes and NKG2D ligands after 48 hours of treatment with 10 µM of ATRA or DMSO. (a-c and e-g) Significance calculated via two-tailed Student’s *t*-test. * p < 0.05, ** p < 0.001. Results are shown for most potent shRBP1 construct of two shRBP1 constructs evaluated.

To address whether *RBP1* is sufficient to reverse the suppressive immune-evasive signature seen in IDHm conditions, stable transduction of a lentivirus expressing RBP1 was employed in IDHm cell lines exposed to retinol, a precursor to ATRA chaperoned by CRBP1. Cell-autonomous markers of NK-mediated immunogenicity, increased NK-mediated cytotoxicity, and a significant increase in several RA-related and Type 1 interferon genes, including *IRF1, IFITM1, STING* and *STAT1* were observed in IDHm astrocytes under *RBP1*-expressing conditions (**Figure 4e-h**). These data demonstrate that *RBP1* is the hallmark of the unique immunomodulatory signature observed in IDHm tumors, and that the absence or presence of *RBP1* determines critical cell-autonomous states that define immunogenic potential in transformed astrocytes.

### ATRA therapy promotes complete tumor regression in a syngeneic glioma model by reprogramming the local and circulating TME in of IDHm murine gliomas

Our *in vitro* human models provided strong rationale to evaluate ATRA’s immunotherapeutic efficacy in genetically faithful, immune competent murine IDHm glioma models. To this end, we utilized a well-validated sleeping-beauty transposon glioma model in C57BL/6 mice (SB-IDHm-R132H)^32^. Tumor-bearing mice were randomized into two treatment groups: 10 mg/kg ATRA every other day and DMSO. ATRA-treated mice demonstrated complete tumor regression by MRI imaging in 5 out of 6 syngeneic IDHm intracranial gliomas (**Figure 5a-d**). Immune phenotyping in this experimental cohort revealed ATRA yielded a >5 fold increase in the number of CD45^+^ TILs per gram of tumor (**Figure 5e**). To understand the immunomodulatory effects driving this increased immune infiltration, we utilized multiparametric flow cytometric phenotyping of tumor infiltrating leukocytes (TILs) from ATRA-treated SB-IDHm tumors. This demonstrated profound ATRA-induced remodeling along several immune lineages in IDHm, but not IDHwt, gliomas (**Figure 5f, Extended Data Figure 4a**). In IDHm gliomas, every immune cell type profiled demonstrated a statistically significant change in upon ATRA treatment. This included the expansion of effector lymphocyte populations (NK, CD8^+^ T cells), induction of antigen presenting cells (M1 macrophages, dendritic cells), and a relative reduction of immunosuppressive cell types (regulatory T cells, monocytic MDSCs, and neutrophilic MDSCs) (**Figure 5f; Supplementary Table 6-7**).

**Figure 5.**
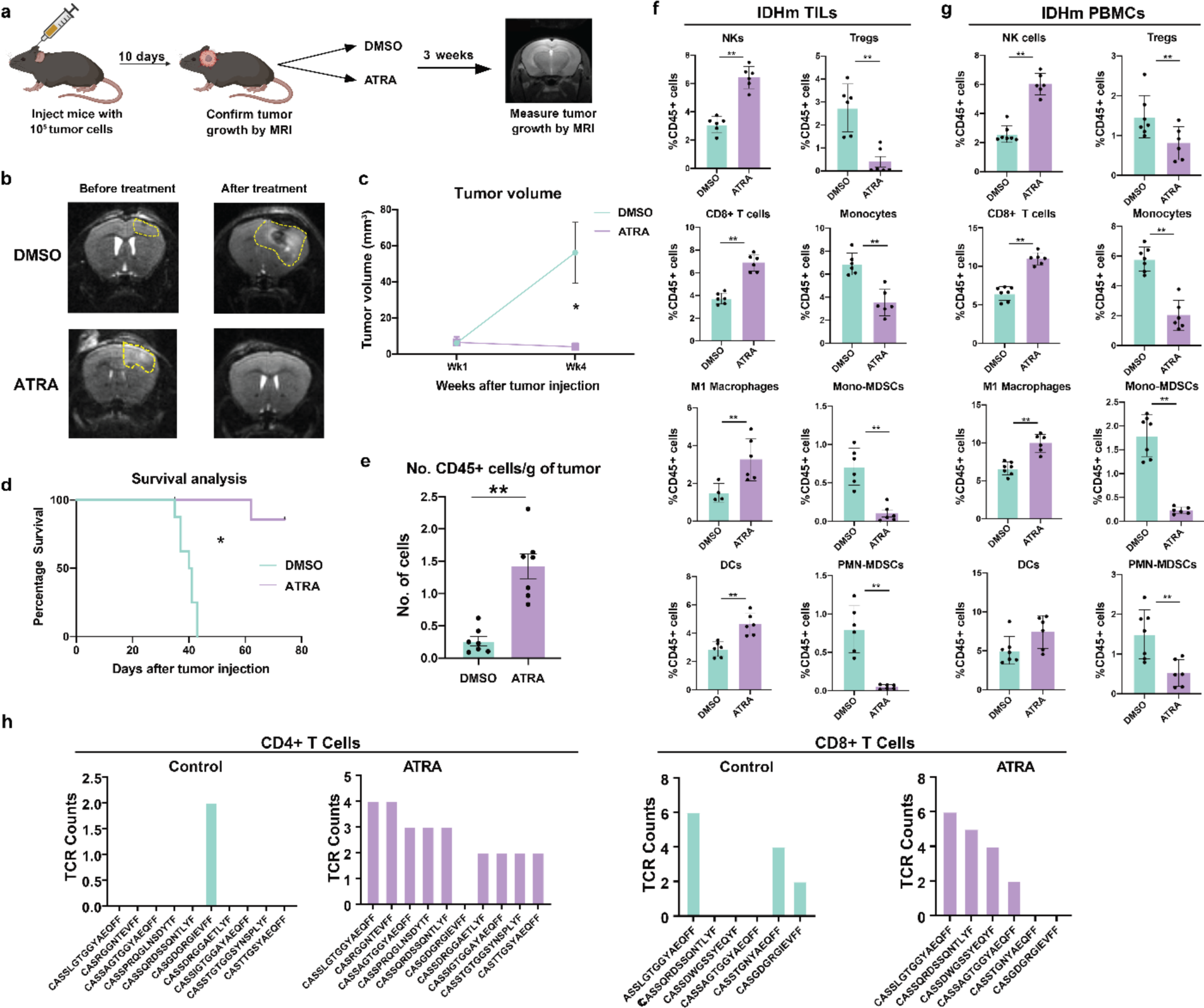
ATRA shows significant therapeutic efficacy in IDH mutant tumor-bearing mice and alters the intratumoral and peripheral immune response. (a) Schematic of experimental design. 6-week old C57BL/6 mice were implanted with 50,000 IDHm tumor cells in the striatum. 10 days after injection, the mice were imaged for tumor presence and volume by MRI. Intraperitoneal ATRA (10mg/kg) or DMSO was administered 3 times a week for 3 weeks. The tumor volume was measured by MRI post-cessation of treatment. (b) Representative MRI images of tumors before and after 3 weeks with DMSO (top panel) or ATRA treatment (bottom panel). (c) Measurement of tumor volume at different timepoints after mice were treated with DMSO or ATRA. (d) Kaplan-Meier survival analysis of IDHm tumor-bearing mice treated with DMSO or ATRA. (e) Quantification of number of immune cells isolated per gram of tumor in DMSO (*n* = 7) and ATRA treated mice (*n* = 7). (f) Flow cytometry analysis of tumor infiltrating leukocytes (TILs) from IDHm tumor-bearing mice treated with DMSO or ATRA. (*n* = 6) (g) Flow cytometry analysis of peripheral blood mononuclear cells (PBMCs) from IDHm tumor-bearing mice treated with DMSO or ATRA. (d-g) Significance calculated via two-tailed Student’s *t*-test. * p < 0.05, ** p < 0.01. (h) T cell receptor (TCR-beta) clonal analysis of CD4^+^ and CD8^+^ tumor infiltrating T cells in ATRA-treated and DMSO-treated SB-IDHm mice. (*n* = 3 pooled samples each)

To gain a comprehensive understanding of ATRA’s systemic immunomodulatory properties in IDHm gliomas, we performed analogous multiparametric spectral flow cytometry on peripheral blood mononuclear cells (PBMCs). Surprisingly, we observed potent pro-inflammatory effects in the circulating immune landscape of IDHm glioma, but no significant differences in the composition of PBMCs between DMSO and ATRA-treated IDHwt gliomas (**Figure 4g, Extended Data Figure 4b**). Indeed, flow cytometric analysis of circulating immune cells (CICs) in ATRA-treated SB-IDHm-bearing glioma mirrored ATRA’s effects on IDHm TILs, resulting in similar immune alterations. We observed significantly increased NK cells, CD8^+^ T cells and M1 macrophages. Conversely, circulating Tregs were reduced by almost 50% and MDSCs were almost completely abrogated with greater than 90% reduction (**Figure 5g**). Interestingly, when comparing tissue-infiltrating and circulating immune landscapes in ATRA-treated IDHm mice, we observed that immune cells were altered in a manner consistent with their expected topographic location. For example, suppressive monocytic cells, which are predominantly found in the systemic circulation, were proportionally decreased to a larger extent in PBMCs than TILs upon ATRA treatment in SB-IDHm tumors. Conversely, regulatory T cells (Tregs), which are primarily found in solid tumor tissue, were decreased to a larger extent in TILs than PBMCs following ATRA treatment.

Using a second murine glioma model, immunocompromised BALB/c nude mice implanted with subcutaneous xenografts (**Methods**), we conducted a limited experimental cohort intended to define whether ATRA’s immunotherapeutic effects where more dependent on innate or adaptive immunologic states (note that BALB/c possess NK but not T cells). These data showed that ATRA dramatically slowed, but did not eradicate, human xenograft IDHm gliomas and similarly altered the myeloid landscape to induce anti-tumor M1-like myeloid phenotypes (**Extended Data Figure 5a-f**). Even in xenograft models, no repression of IDHwt tumor was observed (**Extended Data Figure 5a**). Importantly, the presence of NK cells in BALB/c xenograft models allowed us to recapitulate our *ex vivo* NK cytotoxicity data (**Extended Data Figure 5f-h**), showing that ATRA-mediated tumor regression is dependent on NK cells vitality *in vivo*. NK cell blockade using anti-NK1.1 antibody inhibition rescued tumor growth, albeit with more modest immunomodulatory changes in the TME compared with immune competent animals (**Extended Data Figure 6a-c**). These results suggest that innate and adaptive immune states are required for eradication of IDHm gliomas by ATRA and that NK cells execute important immune cytotoxic effects initiated by ATRA in IDHm gliomas.

An important translational feature of our study is the evaluation of multiple other CNS-penetrant retinoids to determine their efficacy against both IDHwt and IDHm gliomas. It is notable that no other retinoid produced comparable tumor stasis/cytotoxicity compared to ATRA-treated mice, including 13-cis RA or bexarotene (**Extended Data Figure 7a**). Moreover, ATRA induced higher levels of NKG2DL ligands expression *in vivo* with higher relative increases of NK-mediated cytotoxicity *ex vivo* in ATRA-treated mice. The extent of anti-tumor immune infiltration was also markedly higher for ATRA than bexarotene or 13-cis RA (**Extended Data Figure 7b-d**). Taken together, it is apparent that ATRA-mediated anti-tumor effects in IDHm gliomas are characterized by robust immunomodulation involving regulation of circulating and local tumor-infiltrating immune compartments, and that ATRA is likely superior to other CNS penetrant retinoids as an IDH genotype-specific immunotherapeutic monotherapy.

### CD45^+^ immune single-cell sequencing reveals ATRA-induced shifts in myeloid states and increased T cell clonal diversity

To assess the immune cell transcriptomes induced by ATRA in the IDHm glioma TME, we performed pan-CD45^+^ single-cell RNA sequencing (scRNA-seq) in SB-IDHm mice treated with ATRA or DMSO control. We identified diverse myeloid and lymphoid populations, and consistent with our flow cytometric analyses, ATRA radically altered the immune cell composition of TILs in IDHm glioma (**Extended Data Figure 8a-d**). Primarily, we observed differences in the myeloid composition of the ATRA-treated tumor. Namely, M2 macrophages, expressing *Arg1*, *Tgfbi*, and other tumor permissive markers were significantly reduced upon ATRA treatment. Additionally, a population of microglia emerged with high expression of canonical microglial markers *P2ry12*, *Tmem*119 along with increased expression of inflammatory genes such as *Il1a* (**Extended Data Figure 8c-e**), implying homeostatic control of myeloid lineage maturation/activation by ATRA. Examination of cell type compositional changes induced by ATRA also revealed modest increases in the proportion CD4^+^ and CD8^+^ T cells compared to DMSO controls (**Extended Data Figure 8e**). These data, which are consistent with our flow cytometric analyses (**Extended Data Figure 8f**), support that the effect of RA pathway deficiency in IDHm glioma is immunosuppressive and that ATRA efficiently induces anti-tumor myeloid lineages, which likely contribute to its therapeutic efficacy in IDHm glioma.

Since ATRA induced complete therapeutic responses in our syngeneic IDHm glioma model, we hypothesized that adaptive immune responses would be enhanced in IDHm gliomas treated with ATRA. To gain further insight into the effect of ATRA treatment on T-cells in the IDHm TME, we performed T-cell receptor beta (TCRβ) sequencing in conjunction with our CD45^+^ scRNA-seq experiments. Notably, ATRA treatment markedly increased the diversity of clonally expanded CD4^+^ T cell receptor sequences, suggesting more effective adaptive T-cell responses (**Figure 5h**).

### CCL2 chemokine mediates ATRA’s positive effects on immune cell infiltration in IDHm glioma

Having observed ATRA’s potent genotype-specific effects on the immune TME of IDHm glioma *in vivo*, we investigated potential chemotactic factors associated with ATRA’s immunomodulatory function. To this end, we first performed a transwell assay with IDHm or IDHwt astrocytes and healthy donor PBMCs and observed increased migration of NK cells, CD8^+^ T cells and M1 macrophages under ATRA-treated IDHm conditions compared with DMSO-treated controls. Conversely, migration of Tregs, monocytes and M2 macrophages were significantly decreased (**Figure 6a**). These results further emphasize ATRA’s genotype specificity and raised the possibility that soluble chemotactic factors may be mediating this phenomenon. Next, we used a 31-protein ELISA-based chemokine array to identity the ligands mediating these interactions. Interestingly, we observed CCL2 and CXCL12 were the only significantly altered chemokines in ATRA-treated IDHm cells compared with DMSO controls. CCL2 protein production was increased solely in ATRA-treated IDHm cells, whereas CXCL12 was significantly reduced (**Figure 6b**). Quantitative PCR analysis of ATRA or DMSO-treated IDHm astrocytes were concordant with our ELISA data, as ATRA-treated IDHm tumors displayed increased transcription of *CCL2* and reduction of both splice isoforms of *CXCL12* (**Figure 6c**). Repeating the experiment in the presence of a CCL2 blocking antibody confirmed the significance of CCL2 by confirming that CCL2 blockade eradicated ATRA-induced NK and T cell chemotaxis in IDHm, but not IDHwt conditions (**Figure 6d**).

**Figure 6.**
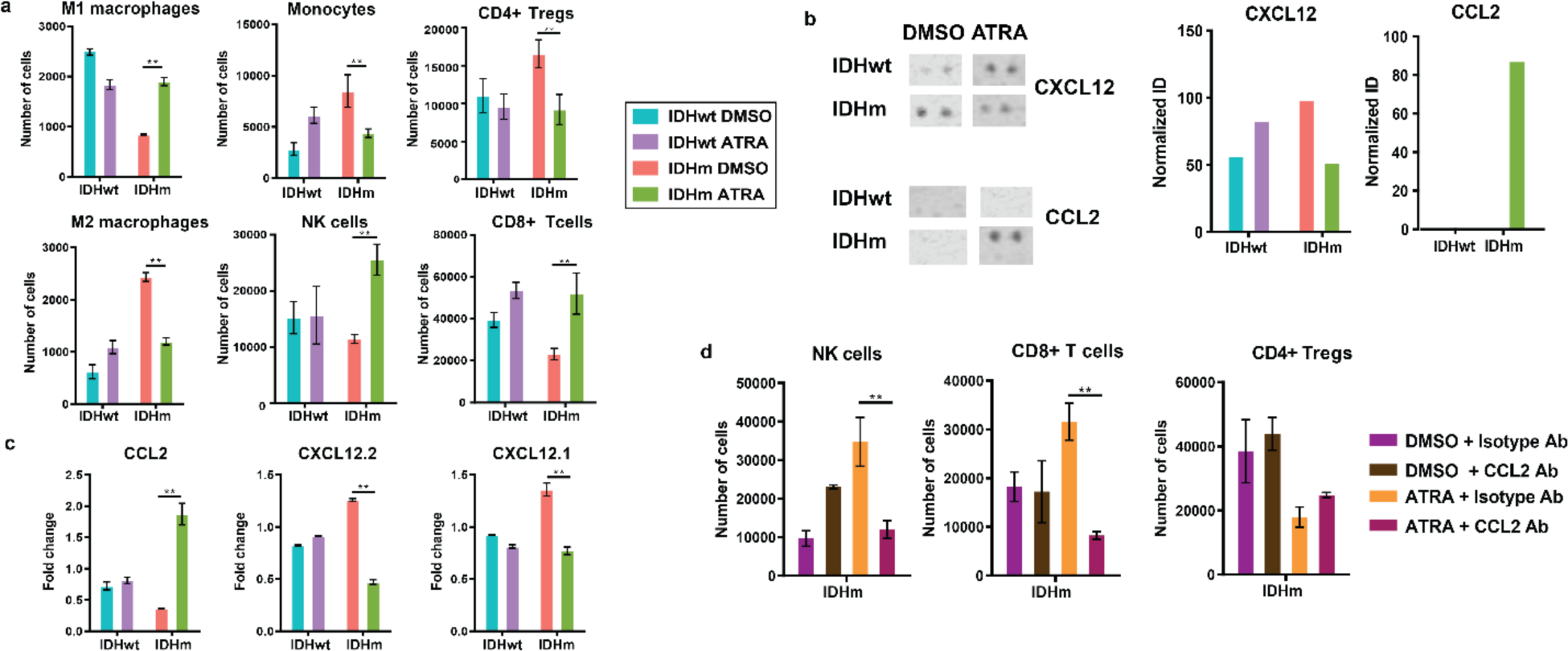
Induction of NK cell chemotaxis by ATRA is dependent on tumor cell-derived CCL2. (a) IDHm and IDHwt GSCs were treated with either DMSO or ATRA for 48 hours in a transwell assay (*n* = 3). Immune cells were harvested from the lower chamber and their quantities are represented in the bar graphs. (b) Left panel: dot blots from chemokine array for CXCL2 and CCL2 performed on supernatants of media from the IDHm and IDHwt tumor cells treated with ATRA or DMSO for 48 hours from the transwell assay. Right panel: Integrated density analysis of the dot blots. The density was normalized to density of control blots (gp100). (c) Gene expression analysis of chemokine transcripts *CCL2, CXCL12.2*, and *CXCL12.1* in primary glioma cells used in the transwell assay after 48 hours of treatment with ATRA or DMSO, as assessed by qPCR (*n = 3*). (d) Transwell assay as in (a) in which cells were cultured with DMSO or ATRA for 48 hours with the presence of either an isotype antibody or an anti-CCL2 Ab. Number of immune cells on the bottom wells of the transwell assay after 48 hours are shown. (a, c-d) Significance calculated via two-tailed Student’s *t*-test. ** p < 0.005.

### ATRA promotes objective tumor regression in human subjects with IDH mutant glioma

Although large phase III trials of ATRA in high-grade gliomas have yet to be conducted, sufficient responses were seen in a subset of patients to justify its use as a second or third-line agent in recurrent gliomas. Consistent with this practice, eight (8) patients with recurrent IDH1/2 mutated gliomas were treated with retinoic acid salvage therapy (ATRA or 13-cisRA) between 2012 and 2018 at the University of Pittsburgh Medical Center (**Supplementary Table 5**). Within this cohort of patients, 4 of 5 IDHm patients receiving ATRA (45mg/kg/m^2^) experienced objective radiographic responses by RANO criteria using the ABC/2 volumetric criteria (**Figure 7a-b**). Notably, ATRA induced statistically significant tumor regression compared to 13-cis RA (**Figure 7a-b**), and all IDHwt patients progressed within 3 months of ATRA initiation (not shown). Even in this small sample size, overall survival in the ATRA-treated patients was significantly longer than 13-cis RA (p<0.05), while progression free survival (PFS) was longer but statistically non-significant (**Figure 7c**). These data, though retrospective, generates significant excitement since no known immunotherapeutic modality produces consistent regression of contrast-enhancing disease in recurrent IDHm glioma.

**Figure 7.**
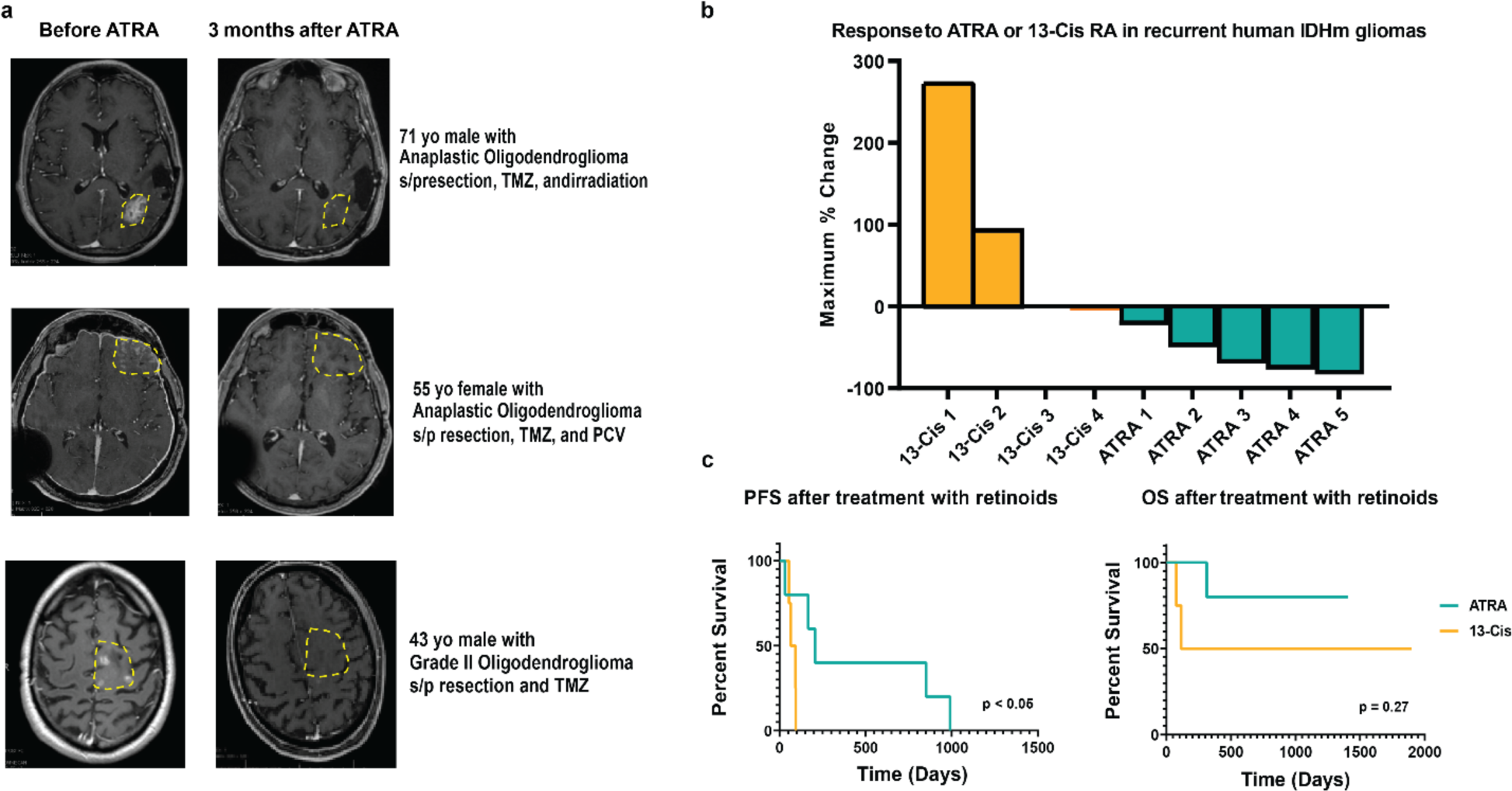
ATRA induces objective tumor response in IDHm glioma patients. (a) Brain MRI of three patients exhibiting significant objective responses to ATRA treatment (delivered at a dose of 45mg/m^2^). (b) Tumor volume calculated using the ABC/2 method^41^ on contrast enhancing lesions on MRI images from patients treated with 13-cis RA or ATRA. (c) Kaplan-Meier survival plot of progression-free survival (PFS) (left) and overall survival (OS) (right) comparing patients receiving 13-cis RA (*n* = 4, 60mg/m^2^) or ATRA (*n* = 5, 45mg/m^2^).

## DISCUSSION

This study identifies potential genotype-specific therapeutic efficacy for all-trans retinoic acid (ATRA) in IDHm gliomas and elucidates unique mechanistic properties that underlie immune-mediated tumor regression in IDHm gliomas. Our study was driven by two salient observations in IDHm gliomas. First, intracellular production of all-trans retinoic acid (ATRA) is nearly absent in IDHm tumor cells compared to IDHwt tumor cells or normal astrocytes. Second, reconstitution of RA signaling through administration of ATRA induces a profound transformation of the IDHm glioma immune microenvironment from immunosuppressive to more inflammatory. A compelling feature of ATRA in IDHm gliomas is its distinct ability to induce chemotaxis of anti-tumor TILs, including cytotoxic T and NK cells, while inducing the expression of activating NK cell ligands in IDHm neoplastic cells. This finding suggests the reversal of repressive immune homeostatic signals maintained by biological mechanisms downstream of oncogenic IDH mutations.

Retinoic acid is a powerful developmental morphogen whose transcriptional output is generally associated with the induction of cellular differentiation programs^33^. Although promulgation of tumor cell differentiation by RA is widely accepted as its primary anti-neoplastic mechanism, it is now abundantly evident that RA maintains immunogenicity in neoplastic states where its activity was thought to be entirely dependent on differentiation, including promyelocytic leukemia (APML)^34,35^ (**Supplementary Discussion**). Overall, robust anti-tumor responses in the IDHm microenvironment by increasing NK cells, decreasing MDSC infiltration, inducing NKG2D stress ligands and NK cell-mediated cytotoxicity, resulting in reduced tumor growth in murines. MDSCs have been consistently identified as prominent drivers of immune suppression in the tumor microenvironment of solid malignancies and their presence is consistently associated with negative responses to immunotherapy ^36,37^. Low MDSC levels in peripheral blood correlate with improved responses to immune checkpoint inhibition in human malignancies^37^. The role of ATRA as a negative regulator of both tissue-infiltrating and peripheral blood MDSCs supports further investigation of multimodal immunotherapy strategies with ATRA and checkpoint inhibitors warrants.

Crucially for clinical translation, we identified differential efficacy of different retinoids. This may be due to select transcriptional programs induced uniquely by ATRA. While retinoid-mediated transcription is broadly determined by ligands binding to RAR/RXR transcription factor dimers, it is evident that different retinoids produce significantly variable transcriptional programs depending on RAR/RXR receptor subtype specificity, tissue variability, and the status of transcriptional co-repressors and co-activators. Our evaluation of blood-brain barrier and white matter-penetrant retinoids reveal underlying differences that may be due to RAR/RXR receptor specificity. For instance, ATRA is a pan-RAR ligand; 13-cis RA binds both RARs and RXRs; and bexarotene is a pan-RXR ligand^38,39^. These data suggest that future evaluations of novel retinoids in IDHm gliomas should skew towards those with strong RAR binding.

The present study establishes important mechanistic characteristics underlying therapeutic efficacy of retinoids in IDH mutant gliomas. We highlight an important and understudied immunotherapeutic role of ATRA and discover key genotype-restricted vulnerabilities to retinoid therapy in glioma. Based on the thorough pre-clinical data established, we inaugurated an ongoing clinical study of ATRA in conjunction with PD-1 inhibition to determine whether retinoid therapy is a viable option for the management of recurrent human IDH mutant gliomas.^40^

## Supporting information

Supplementary Discussion

## Acknowledgements

This work was supported by NIH grant P30CA047904, the Karp Family Foundation, the United Initiative to Cure Brain Cancer, Hillman Post-doctoral Fellowship for Innovative Cancer Research, and The University of Pittsburgh holds a Physician-Scientist Institutional Award from the Burroughs Wellcome Fund. We would like to thank Dr. Maria G. Castro who generously provided orthoptic models of glioma, and Dr. Timothy Chan who provided IDH wildtype and mutant glioma cell lines.

## Author Contributions

Conceptualization, A.R., X.Z., and N.M.A.; methodology, A.R., X.Z., P.S., and N.M.A; experimental design, A.R., X.Z., P.S., R.P., J.B., A.N.M., E.H., and N.M.A.; conduction of experiments, A.R., X.Z., Y.K., W.J.K., E.J., J.A., R.P., K.K., and C.C.; resources, N.M.A.; bioinformatics analysis, A.R.C., J.H.S., J.X.; writing, A.R., S.A., E.H., J.H.S., A.C.T., J.P., E.A., A.A., A.L., R.F., A.D., E.C.H., S.A, A.V., P.O.Z, S.J.B., T.C.B., D.A.V., and N.M.A.; funding acquisition, N.M.A.; supervision, N.M.A.

## Competing Interests

The authors declare the following competing interests: N.M.A. and A.R. have a patent on Retinoid composition and use for IDH-mutant cancers (US Patent 10821091).

## METHODS

### Lead Contact

Further information and requests for resources and reagents should be directed to and will be fulfilled by the lead contact, Nduka M. Amankulor

### Materials availability

All unique and stable reagents generated in this study of which sufficient quantities exists are available from the Lead Contact with a completed Materials Transfer Agreement.

### Data and code availability

Data that was generated during the study have been deposited in Gene Expression Omnibus (GEO) with the accession number GSE252527 (bulk RNA-seq of IDHwt and IDHm astrocyte cell lines treated with ATRA) and GSE172387 (scRNA-seq of intratumoral CD45^+^ sorted cells from ATRA-treated and control mice). Detailed scripts and parameters used for each step of the analysis provided by reasonable request to the authors.

### Cell lines and Cell Culture

Immortalized human syngeneic astrocyte cell lines with retrovirally transfected IDH1-R132H construct, IDHwt construct and mock-transfected with a retroviral vector were obtained from Dr. Tim Chan’s group (MSKCC). The cells were cultured in Dulbecco’s Modified Essential Medium (DMEM) supplemented with 10% FBS, 2 mM L-glutamine, 100 U/ml penicillin, and 100 μg/ml streptomycin (Complete Medium, (CM). All reagents were obtained from HyClone™ (GE Healthcare). The cells were maintained in a humidified incubator at 37°C and 5% CO_2_. Medium was replaced every 2 days and the cells were split when they were ∼80% confluent. All cell lines were negative for mycoplasma infections. For glioma stem cell culture, TS603 and TS667 were obtained from Dr. Tim Chan’s group^8^. These cells have been extensively studied and described in previous reports^42^. The cells were maintained in Neurocult™ basal proliferation medium (Stemcell Technologies, Vancouver) containing 10ng/ml EGF, 10ng/ml FGF and 0.0002% Heparin (Stemcell Technologies, Canada).

### Glioma cell culture

Resected tumors from Glioma patients (both IDHwt and IDHm) were mechanically dissociated to form a single cell suspension. Contaminating RBCs were removed using an RBC lysis buffer (eBioscience) and cells were propagated as neural stem cells in Neurocult™ NS-A proliferation kit supplemented with 20 ng/mL Human Recombinant EGF, 10 ng/mL Human Recombinant bFGF) and 2 μg/mL Heparin Solution (Stemcell Technologies Canada Inc.). For a subset of experiments neurospheres were moved to Complete Medium (CM) to propagate differentiation into adherent cell lines. All tumors were genotyped using GlioSeq™ technology to obtain IDH mutation status.

### Transwell Immune Cell Migration Assay

We modeled the immune tumor microenvironment of gliomas *in vitro* using a transwell assay with IDHm or IDHwt GSCs treated with ATRA or DMSO and plated on the bottom chamber of a transwell. PBMCs were placed on the top chamber at a ratio of 1:10 GCS to PBMC. Immune cells migrating to the bottom chamber were collected and analyzed using flow cytometry after 48 hours.

### ELISA based Chemokine Array

IDHwt and IDHm GSCs were cultured for 48 hours with ATRA or DMSO and the supernatants were collected and used to assess changes in chemokine expression using a chemokine array kit.

### Mice

#### Immunocompetent mouse model

We used the Sleeping Beauty transposase model to generate IDHm glioma neurospheres as previously described^43^. The neurospheres were maintained in DMEM-F-12 medium containing 20ng/ml of bEGF, bFGF and 1ug/ml Normocin. 50,000 cells in 3µl volume were injected orthotopically into the striatum of 6-week old female C57BL/6 mice. The mice were imaged for tumor growth by MRI pre-treatment and were sacrificed when they showed signs of CNS pathology and the tumors were harvested as follows. Tumors were sharply dissected away from surrounding brain tissue based on appearance and tactile feel. They were then chopped into 1-2mm pieces and incubated in Accutase (Corning) for 10 minutes at 37°C. The suspension was then sequentially passed through 25ml, 10ml, and 5ml pipettes. Further dissociation was done by passing the suspension through 18G needle 3 times and then through a 100-micron strainer. The remaining suspension was then spun down and resuspended in 1ml of RBC lysis buffer for 10 minutes in room temperature after which the cells are spun down and resuspended in a sucrose gradient. For ATRA treatment, 10 mg/kg ATRA was administered to mice intra-peritoneally every other day until termination of the experiment. Control mice were treated with DMSO on the same schedule.

#### Xenograft mouse model

6-8 week old BALB/c nude mice (CAnN.Cg-*Foxn1nu*/Crl) were purchased from Charles River Laboratories. Mice were injected with 10^6^ IDHwt or IDHm primary glioma cells subcutaneously in the right flank. When the tumors reached about 25mm^2^ (typically around 10 days post-tumor inoculation), mice were randomized into treatment groups and ATRA was initiated. 10 mg/kg ATRA was administered to mice intra-peritoneally every alternate day until termination of the experiment. Control mice were treated with DMSO on the same schedule. Animals were monitored for signs of drug toxicity, including weight loss, lethargy, skin lesions, and diarrhea. Tumor measurements were obtained every 3 days using digital vernier calipers. Mice were sacrificed when tumor volumes reached ∼400 mm^3^. For NK depletion experiments, 6-week old B6 nude mice (B6.Cg-Foxn1nu/J) were purchased from The Jackson Laboratory. NK depletion was performed by injecting mice with 200μg of anti-NK1.1 antibody intra-peritoneally concurrently with ATRA initiation. Mice injected with 200μg of isotype antibody were used as controls. Peripheral blood was collected via tail vein for quantification of circulating NK cells by flow cytometry.

### All-trans retinoic acid (ATRA) and 2HG-inhibitor treatment of astrocytes and primary glioma cells

ATRA was purchased from Sigma Aldrich and reconstituted in DMSO. ATRA was added at a 1µM final concentration to cell culture medium for 48 hours. For differentiation experiments, IDHwt or IDHm neutrospheres were treated with ATRA for 14 days continuously. After treatment, the cells were either lysed in Trizol for qPCR analysis or were used in NK cytotoxicity assays. The small molecule inhibitor of 2HG, AGI-5198, was used in selected experiments.

### ATRA Bioanalysis

The liquid chromatography system consisted of an Agilent (Palo Alto, CA, USA) 1200 SL autosampler and binary pump, a Phenomenex (Torrance, CA USA) Kinetex C18 (2.6 μm, 50 x 2.1 mm) column, and an isocratic mobile phase. Mobile phase solvent A was 0.1% (*v/v*) formic acid in acetonitrile, and mobile phase solvent B was 0.1 % (*v/v)* formic acid in water. Solvent A was pumped at 57% and solvent B was pumped at 43% with a total flow rate of 0.5 mL/min. The run time was 12 min.

Mass spectrometric detection was carried out using an ABI SCIEX (Concord, ON, Canada) 4000Q hybrid linear ion trap tandem mass spectrometer with electrospray ionization in positive multiple reaction monitoring (MRM) mode. The settings of the mass spectrometer were as follows: curtain gas 40, CAD 10, Ion transfer voltage 5000 V, probe temperature 500°C, GS1 40, GS2 40, declustering potential 50 V, entrance potential 5 V collision energy 20 V, and collision cell exit potential 10 V. The MRM *m/z* transitions monitored were: 301.5>123.5 for ATRA and 306.5>127.5 for [D_5_]-ATRA (internal standard). The LC system and mass spectrometer were operated as previously reported with 1/y^2^ weighted linear regression.

All sample handling occurred under sodium light. The matrix for standard curve samples and dilution of cell samples was 10% human plasma in distilled water. A volume of 10μl of internal standard (1 µg/mL [D_5_]-ATRA) was added to each 100μl of standard or sample cell/plasma matrix. Next, 1ml of acetonitrile was added followed by vortexing for 1 min on a Vortex Genie-2 set at 10 (Model G-560 Scientific Industries, Bohemia, NY, USA). Samples were centrifuged at 12,000 × *g* at room temperature for 5 min. Supernatants were transferred to 12 mm × 75 mm borosilicate glass tubes and evaporated to dryness under a stream of nitrogen at 37°C. Dried residues were re-dissolved in 100μl of acetonitrile:water:formic acid (60:40:0.1, *v/v/v*). After brief vortexing, the supernatants were transferred to autosampler vials, followed by injection of 10μl into the LC-MS/MS system. The assay was linear from 1–1,000 ng/ml. Accuracy (89.8-109.0%) and precision (<11.7%CV) were acceptable^44^.

### *RBP1* silencing and overexpression studies

For *RBP1* silencing, lentiviral constructs bearing *RBP1* shRNA (Santa Cruz biotechnologies), or scrambled shRNA sequences (mock) were transfected in IDHwt glioma cells (Santa Cruz biotechnologies). *RBP1*-silenced clones were selected in the presence of 10ug/ml Puromycin in the medium. Mock transfection of scrambled shRNA lentiviral vectors was used as controls, and clones expressing the construct were selected as above.

For *RBP1* overexpression studies, IDHm glioma cells were transfected with *RBP1* lentiviral vectors (Origene Inc) expressing the full *RBP1* cDNA. The cells were maintained in CM containing 10ug/ml puromycin to select for lentivirus-transfected cells. Cells transfected with an empty lentiviral vector were used as controls.

### Isolation of NK cells

#### From donor PBMCs

Peripheral blood samples were collected in preservative-free heparin tubes (10 U/ml) and layered into an equal volume of Ficoll-Hypaque density gradient solution (Amersham Pharmacia Biotech Ltd., Little Chalfont, UK) and centrifuged at 2250 rpm at 20°C. The mononuclear cells (PBMCs) were collected and washed twice with PBS (Hyclone™, GE Healthcare). Cell viability was determined by trypan blue exclusion and exceeded 95%. Isolated PBMCs were used for NK cell isolation using CD56 Microbeads (Miltenyi Biotec Inc.) and a MidiMACS separation column, according to the manufacturer’s instructions. Purity was (>95%) as determined by FACS analysis.

#### From mice

The spleens from naïve BALB/c mice were isolated and mechanically dissociated to form single-cell suspensions. RBCs were lysed using RBC lysis buffer (eBioscience). The cells were used for NK isolation using NK cell isolation kit (mouse) (Miltenyi Biotec Inc.) and a MidiMACS separation column, according to the manufacturer’s instructions. Purity was (>95%) as determined by FACS analysis.

### NK-92 cell culture

For some experiments, NK92 cells were used instead of donor NK cells. NK92 cells (ATCC® CRL-2407™) were obtained from ATCC. The cells were grown in Alpha Minimum Essential medium with 2 mM L-glutamine and 1.5g/L sodium bicarbonate, 0.2 mM inositol, 0.1 mM 2mercaptoethanol, 0.02 mM folic acid (all from Sigma Aldrich) 12.5% horse serum and fetal bovine serum (ATCC) to a final concentration of 12.5%. Fresh medium was added every 2-3 days. Once a week, the cells were counted and re-suspended at a final concentration of 5*10^5^ cells/ml in fresh medium.

### NK cytotoxicity assay

The flow cytometric CFSE/7-AAD cytotoxicity assay was performed as previously described^45^ with slight modifications. Astrocytes or glioma cells (5 × 10^5^) were labeled with 500 nM CFSE (from a 1 mM stock solution in dimethyl sulfoxide [Sigma] stored at −20°C) in PBS for 15 min at 37°C. The cells were then washed twice in complete medium and used immediately for the cytotoxicity assay. The CFSE-labeled target cells (25,000 cells) were used at E (effector):T (target) ratios of 1:10. After 6h incubation, the cells were stained with 0.25 μg/ml of 7-AAD and analyzed immediately by flow cytometry.

### Annexin V apoptosis assay

Apoptotic cells were detected using the ThermoFisher Annexin V staining kit following the manufacturer’s instructions including surface and intracellular staining.

### Chemokine assay

The ELISA-based Proteome Profiler Human Chemokine Array Kit (Biotechne #ARY017) was used, which includes 31 different chemokine antibodies, each spotted in duplicate. The assay was performed following the manufacturer’s instructions.

### Flow cytometry for TILs

IDHwt and IDHm tumors were dissociated to form a single cell suspension (SCS). The SCS was then overlaid onto sucrose gradient (Lympholyte-M) and centrifuged at 2400 rpm for 20 minutes. Immune cells at the interphase of the 2 liquids were collected and washed once with PBS. The cells were then stained with appropriate antibodies and acquired on a BD LSR Fortessa flow cytometer. Briefly, cells were washed with PBS and stained with 1μg antibody for 30 minutes at 4°C. After incubation, the cells were washed three times with FACS buffer (PBS + 10% FBS + 0.1% sodium azide), resuspended in a final volume of 200ul of FACS buffer and run through the flow cytometer. Flow cytometry data was analyzed on the FlowJo v9.1 software. Gating strategies are shown in **Supplementary Table 6.** Antibody panel and catalogue information are included in **Supplementary Table 7**.

### Gene expression profiling by RNA-seq

ATRA-treated IDHwt and IDHmut astrocytes were prepared for bulk RNA-sequencing using Illumina’s TruSeq RNA Access Library Preparation protocol following the manufacturer’s instruction. Pooled libraries were then sequenced on Illumina NextSeq 500 platform with single-end 76 bp reads, to a target read count of 40 million reads per sample. Reads were aligned to the human genome (GRCh38) using STAR aligner(v2.7.3a) with default parameters^46^. Gene-count matrices were produced by featureCounts (v2.0.1)^47^. Normalization of RNA sequencing data and differential gene expression analysis was performed using DESeq2^48^. The *rlog* transform was used to compute the principal component analysis and select top variable genes. LogFC values were shrunk using the *ashr* package^49^ as implemented in DESeq2. Gene set enrichment analysis was conducted using the *fgsea* package^50^ with default parameters, using the DESeq2-statistic to rank the gene list. For direct comparisons between two conditions, we used the *contrast* parameter to compare the conditions. To identify IDHm-specific effects of ATRA treatment, we utilized a linear model of the form ∼IDH + Treatment + IDH:Treatment, where IDH:Treatment is the interaction coefficient. Gene sets were obtained from the Molecular Signatures Database (MSigDB)^51^, including the Gene Ontology and Reactome gene sets. Data is accessible from GEO, accession number GSE172055.

### Single-cell RNA sequencing library preparation

Single-cell RNA sequencing was performed on TILs isolated from both ATRA-treated and untreated tumors. Briefly, tumors were resected from mice as described above and live CD45^+^ cells were sorted on a Beckman Coulter MoFlo Astrios High Speed Cell Sorted at the Hillman Cancer Center Flow Cytometry Core. Sorted cells were then count using AO/PI staining on a Nexcelom Cellometer, and single-cell expression libraries were generated using the Chromium Single Cell 5 ‘Reagent (V1 Chemistry) as per the manufacturer’s instructions. Final libraries were assessed for size distribution and concentration, then pooled for sequencing on a Novaseq 6000 at the UPMC Genome Core with the following sequencing parameters: read 1: 28 cycles; read 2: 91 cycles; i7: 8 cycles.

### Single-cell RNA-seq analysis

Sequencing results were demultiplexed using bcl2fastq (Illumina) using a base mask of Y28, I8, and Y91 and allowing for no barcode mismatches in the i7 index. Alignment of demultiplexed reads was next performed using CellRanger (10X Genomcs, v3.1.0) using the mm10 reference genome. Following alignment, the untreated and treated samples were combined into a unified feature barcode expression matrix, and downstream analysis was performed using Seurat (v3.1.2) Briefly, the top 2,000 most variable genes were selected using the FindVariableFeatures function, and principal component analysis was performed using these highly variable genes. Principal components (PCs) were heuristically selected based on an elbow plot for inclusion in downstream analysis (the first 15 PCs were used). Uniform Manifold Approximation Embedding (UMAP) was used for visualization, with the first 15 PCs as input. Cells were clustered using the *FindNeighbors* and *FindClusters* functions. Differentially expressed genes between clustering were identified using a Wilcoxon rank sum test as implemented in the *FindMarkers* function with default parameters. Cells were annotated based on canonical cell type markers (**Figure 7c-d**). Data is accessible from GEO, accession number GSE172387.

### Quantitative Real time RT-PCR

5×10^5^ IDHm and IDHwt astrocytes, as well as patient glioma cells were lysed in Trizol and RNA was extracted from the cells using the Chloroform extraction method. cDNA was transcribed from the RNA using the iScript cDNA synthesis kit (BioRad). Real-time PCR was performed on a CFX96 Touch™ Real-Time PCR Detection System (Bio-Rad) equipped with a 96-well reaction plate. Briefly, 2µl of cDNA was added to 5µl of the 2× SYBR Green PCR mastermix(Thermo Scientific), 800nM of each primer, and water was added to 10µl. The thermal denaturation protocol was run at the end of the PCR to determine the number of products that were present in the reaction. Reactions were typically run in triplicates. The cycle number at which the reaction crossed a threshold (*C*_T_) was determined for each gene and the relative amount of each gene to 18 S rRNA was described using the equation 2^−Δ*C*^_T_ where Δ*C*_T_ = (*C*_TtargetRNA_ − *C*_T18S rRNA_).

### Treatment of human patients with ATRA

Eight patients with IDH1/2 mutated gliomas were started on retinoic acid as a salvage therapy between 2012 and 2018. 13-cis retinoic acid was used prior to 2017, after which all-trans retinoic acid was used (**Supplementary Table 5)**. Four patients received 13-cis retinoic acid at dose of 50mg/m^2^ per day and five patients received all-trans retinoic acid at 80mg daily. One patient had to reduce the dose of all-trans retinoic acid to 50mg daily due to headaches. Retinoic acid therapy was stopped at the time of new radiographic progression. Progression free survival was calculated based on radiographic progression on MRI. Tumor volume was calculated using ABC/2 method^41^ on contrast enhancing lesions. MRI imaging was reviewed by two neurosurgeons, two neuro-oncologists, and a neuroradiologist for evidence of progression.

### Statistical Analysis

All data points are represented as means ± SD, and the two-tailed Student’s *t* test was used to compare mean values between the two groups unless otherwise indicated. One-way or two-way analysis of variance (ANOVA) was used to compare mean values between multiple groups. Statistical analysis was performed using Prism version 8 (GraphPad Software, La Jolla, CA). *P* values < 0.05 were considered statistically significant, unless otherwise noted. Downstream analysis of bulk and single-cell RNA-seq data was conducted in R v4.3.1.

### Study approval

All relevant mouse studies were approved by the University of Pittsburgh IACUC committee, protocol #19085737. Human clinical data collection was approved by University of Pittsburgh IRB, STUDY1810002. No individual patient consent was necessary.

## Extended Data Figures

**Extended Data Figure 1.**
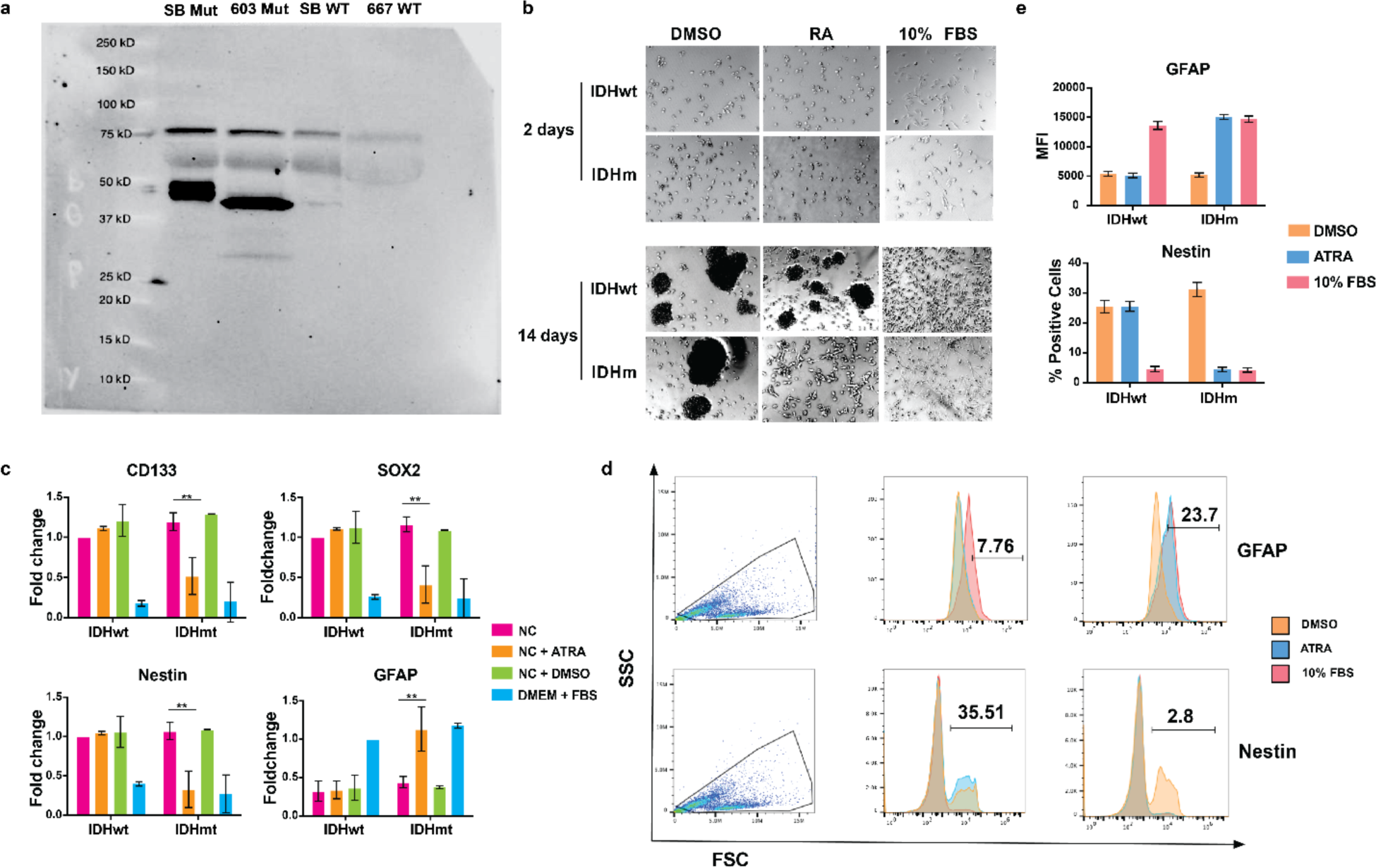
ATRA induces delayed differentiation in IDHm glioma cells. (a) Western blot validation of IDHm glioma cell lines showing expression of the mutant IDH1-R132H protein. (b) Representative images demonstrating morphology of IDHwt and IDHm GSCs after treatment with 1 µM of ATRA, DMSO, or 10% fetal bovine serum (FBS) at 48 hours (top panel) and 14 days (bottom panel) after administration. 10x magnification. Figures are representative of 6 independent fields of view. (c) Gene expression of stem cell and differentiation markers in IDHm and IDHwt GSCs treated with 1µM of ATRA, DMSO, or 10% FBS for 14 days, as assessed by qPCR. ** p = 0.005 as calculated by Student’s *t-*test. (d) Schematic of flow cytometry analysis of ATRA, DMSO, or 10% FBS-treated GSCs for GFAP, and Nestin. (e) Bar graph detailing the flow cytometry data as outlined in (d), showing mean fluorescence intensity (MFI) of GFAP (top panel) and percent of cells positive for Nestin (bottom panel).

**Extended Data Figure 2.**
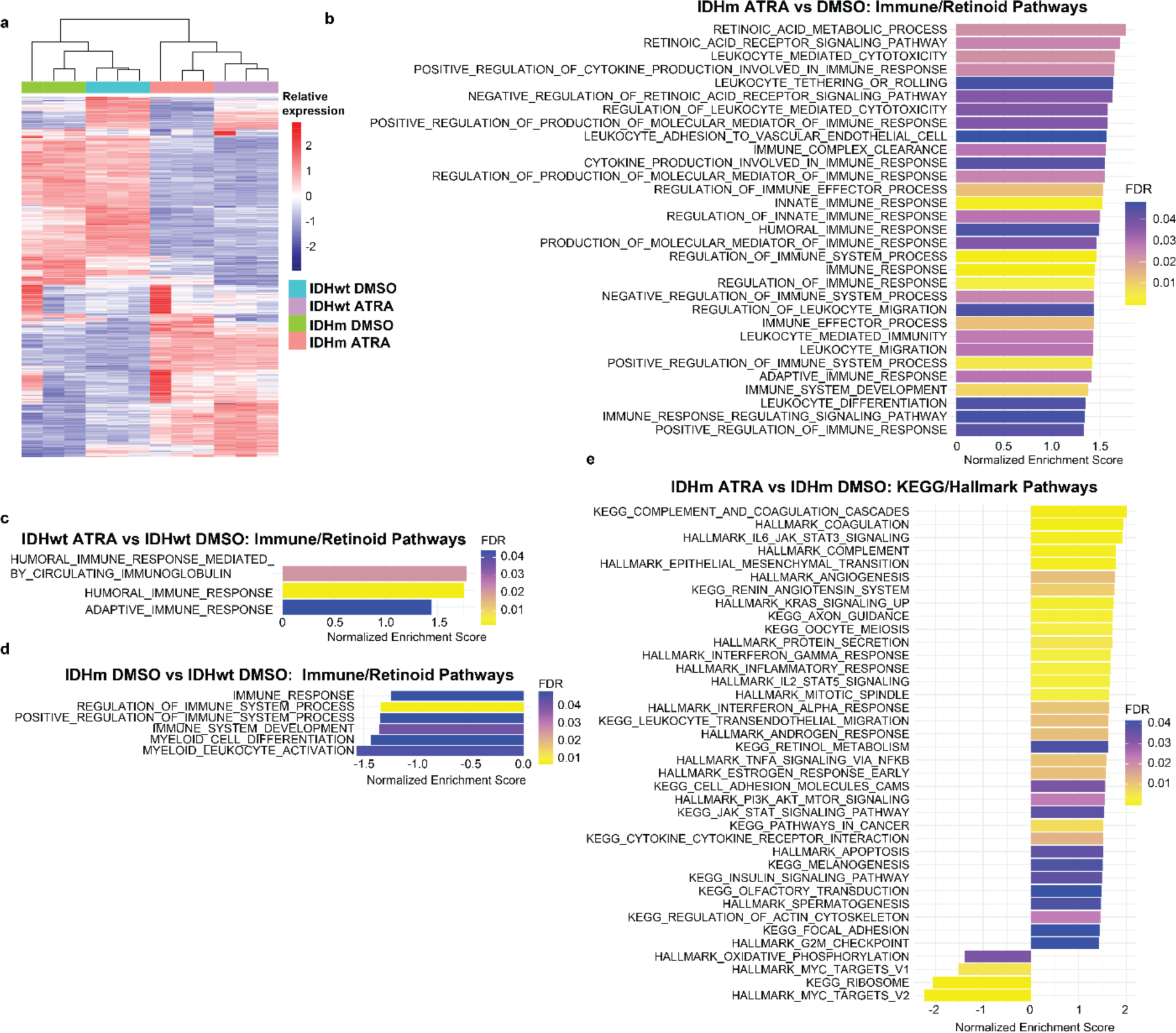
ATRA induces global remodeling of the transcriptome in both IDHm and IDHwt astrocyte cells. (a) Heatmap of relative expression of top 2000 most variable genes across the bulk RNA-seq data of ATRA-treated and DMSO-treated IDHwt and IDHm cell lines. (b) Gene set enrichment analysis (GSEA) of Gene Ontology pathways related to retinoic acid signaling and immune response comparing ATRA-treated versus DMSO-treated IDHm cell lines. (NES, Normalized Enrichment Score; FDR, False Discovery Rate). (c) GSEA of pathways in (b) comparing ATRA-treated versus DMSO-treated IDHwt cell lines. (d) GSEA of pathways in (b) comparing DMSO-treated IDHm and IDHwt cell lines. For (b)-(d), all significant pathways (FDR <0.05) are shown. (e) Gene set enrichment analysis of ATRA-treated versus DMSO-treated IDHm cell lines across pathways from the KEGG and Hallmark databases, showing top upregulated and downregulated pathways.

**Extended Data Figure 3.**
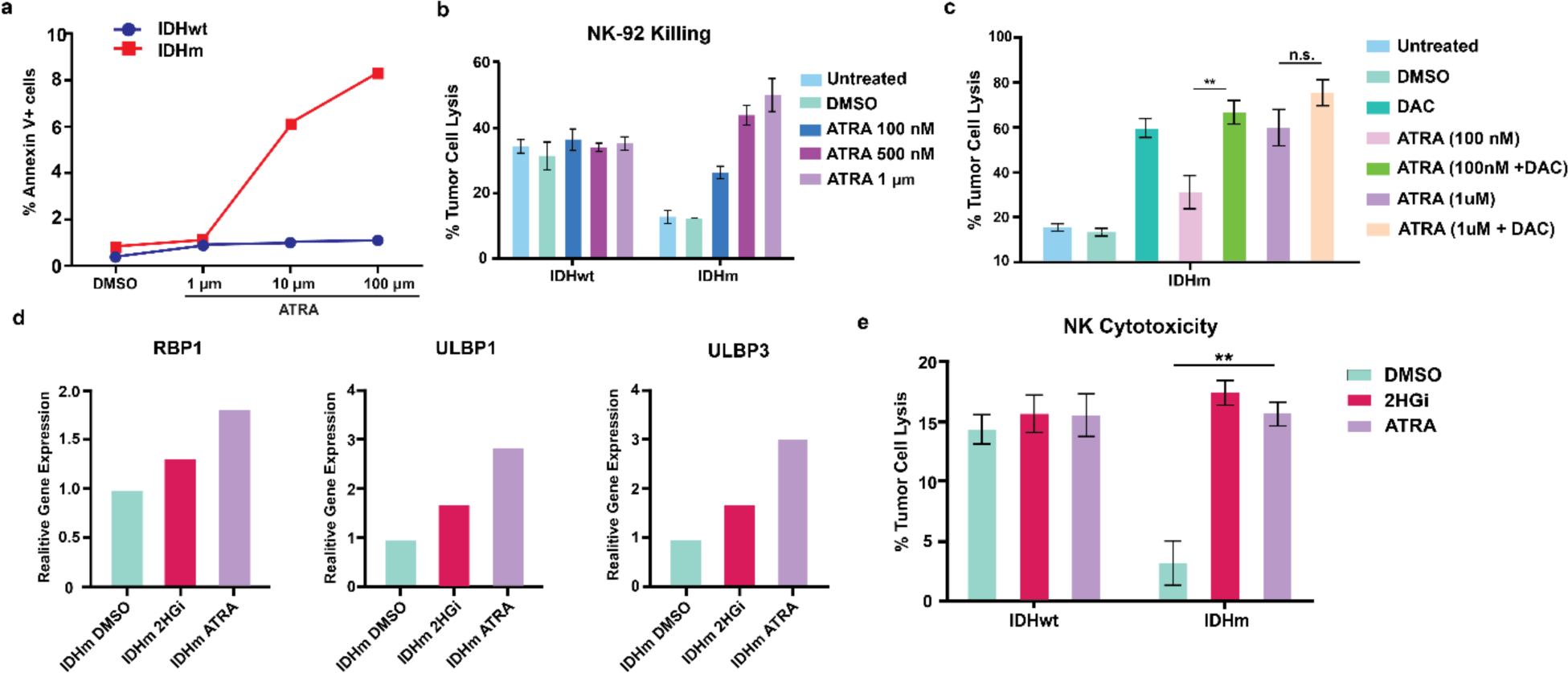
ATRA and 2HG inhibition in IDHm GSCs partially restores *RBP1* and *NKG2DL* expression and increases their susceptibility to NK mediated cytotoxicity. (a) Flow cytometry measurements of percent tumor cell lysis of IDHwt and IDHm GSCs that were cocultured with NK-92 cells and treated with DMSO or increasing concentrations of ATRA. (b) Flow cytometry measurements of percent tumor cell lysis of IDHm GSCs after treatment with DMSO, ATRA, or combination of ATRA and DMSO at two different concentrations as shown in the color legend. * *p* < 0.05, ** *p* < 0.01 calculated by two-tailed Student’s *t*-test. (c) Gene expression of *RBP1*, *ULBP1*, *ULBP3* in IDHm GSCs after treatment with DMSO, ATRA, or 2HG inhibitor (AGI-5198), as assessed by qPCR. (d) Flow cytometry measurements of percent tumor cell lysis as a surrogate for NK-mediated cytotoxicity on IDHm and IDHwt GSCs that were cocultured with NK-92 cells and treated with ATRA, 2HGi, or DMSO. ** *p* = 0.005 as calculated by two-tailed Students *t-* test (e) Surface Annexin V staining on IDHwt and IDHm GSCs after treatment with DMSO or increasing concentrations of ATRA as a surrogate marker for percent of cells undergoing apoptosis.

**Extended Data Figure 4.**
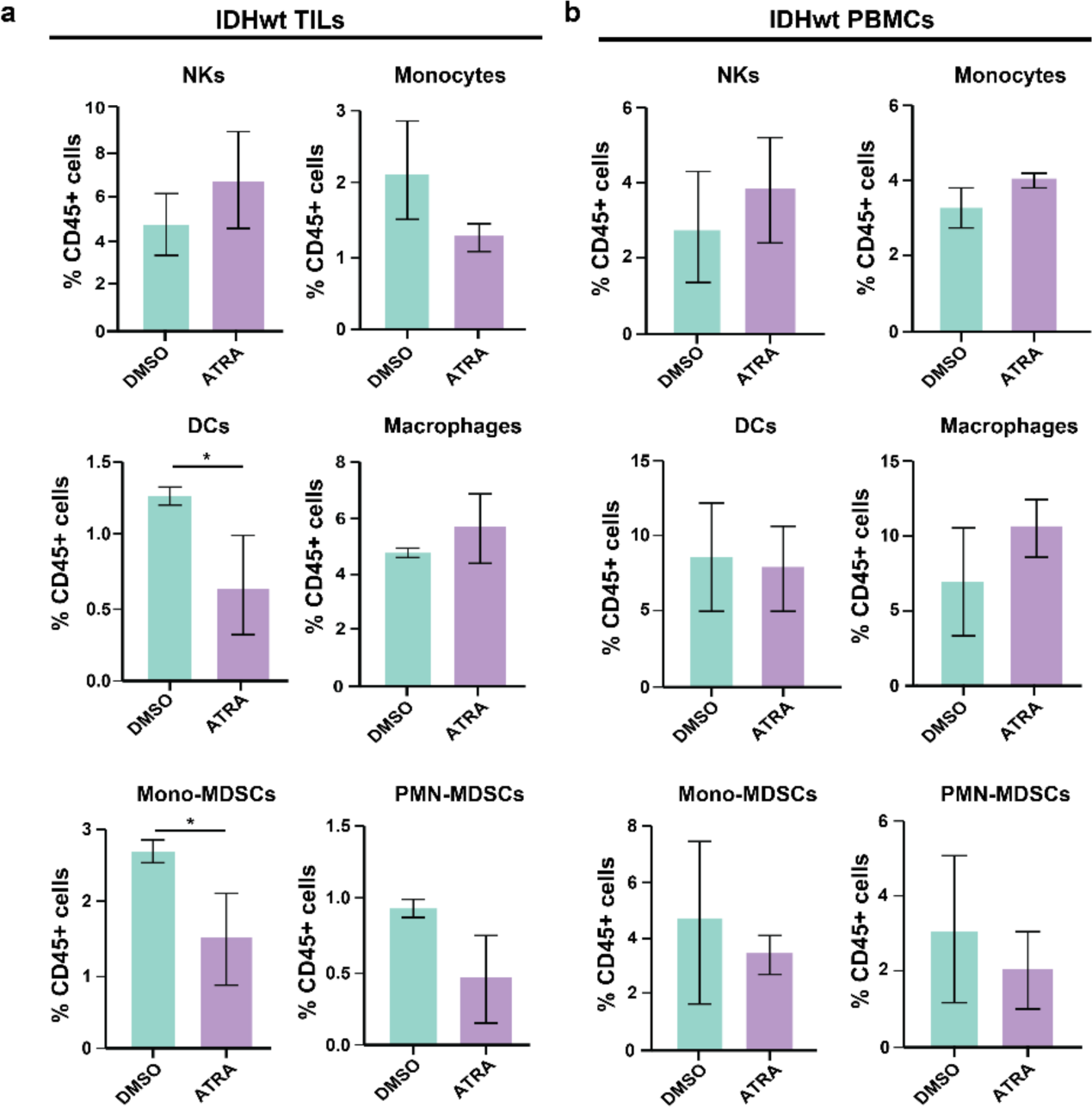
ATRA treatment has limited impact on intratumoral and peripheral immune composition in mice bearing IDHwt tumors. Flow cytometry analysis of (a) TILs and (b) PBMCs from IDHwt tumor-bearing mice treated with 10mg/kg of ATRA or DMSO intraperitoneally 3 times a week for 3 weeks (*n* =3). Significance calculated via two-tailed Student’s *t*-test. * p < 0.05.

**Extended Data Figure 5.**
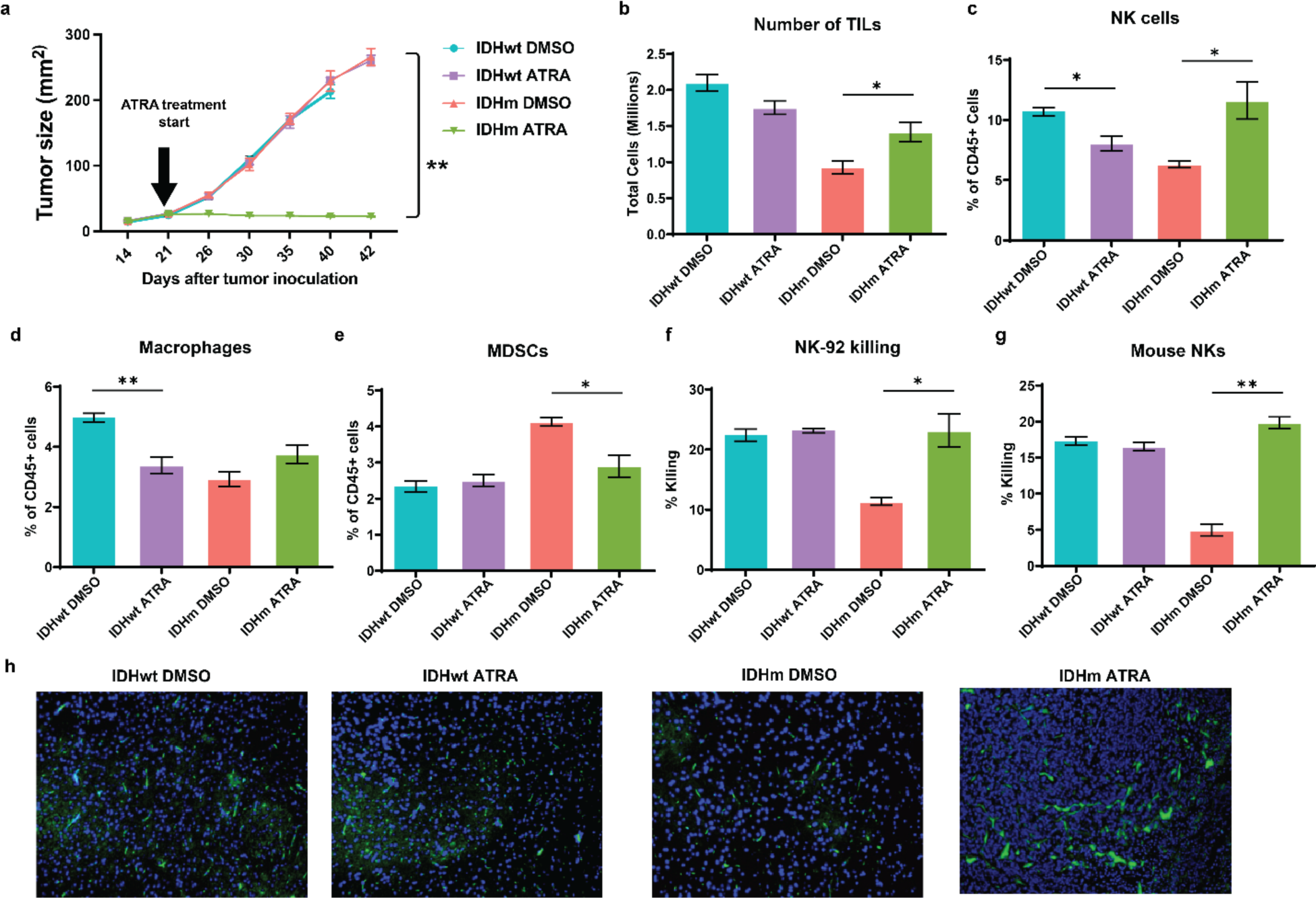
ATRA is effective at inducing tumor stasis in IDHm, but not IDHwt, xenograft tumor-bearing mice *in vivo*. (a) Tumor growth curves of BALB/c nude mice bearing IDHwt or IDHm flank tumors treated with either 10 mg/kg of ATRA or DMSO every alternate day for 21 days. (b) Quantification of immune cell infiltration via flow cytometry in IDHwt or IDHm tumors treated with either 10 mg/kg of ATRA or DMSO every alternate day for 21 days. (c) *Ex vivo* NK cytotoxicity assay using human a NK cell line (NK-92) and tumor cells isolated from ATRA or DMSO-treated mice. (d) *Ex vivo* NK cytotoxicity assay using freshly isolated NK cells from mice. (e) Immunohistochemistry of paraffin-embedded sections of tumor. Nuclei are marked with DAPI (blue) while NK cells are marked with green fluorescent protein. Images shown are representative of 3 independent fields of view. Magnification 10X. (b-g) * *p* = 0.05, ** *p* = 0.005 calculated by two-tailed Student’s *t-*test.

**Extended Data Figure 6.**
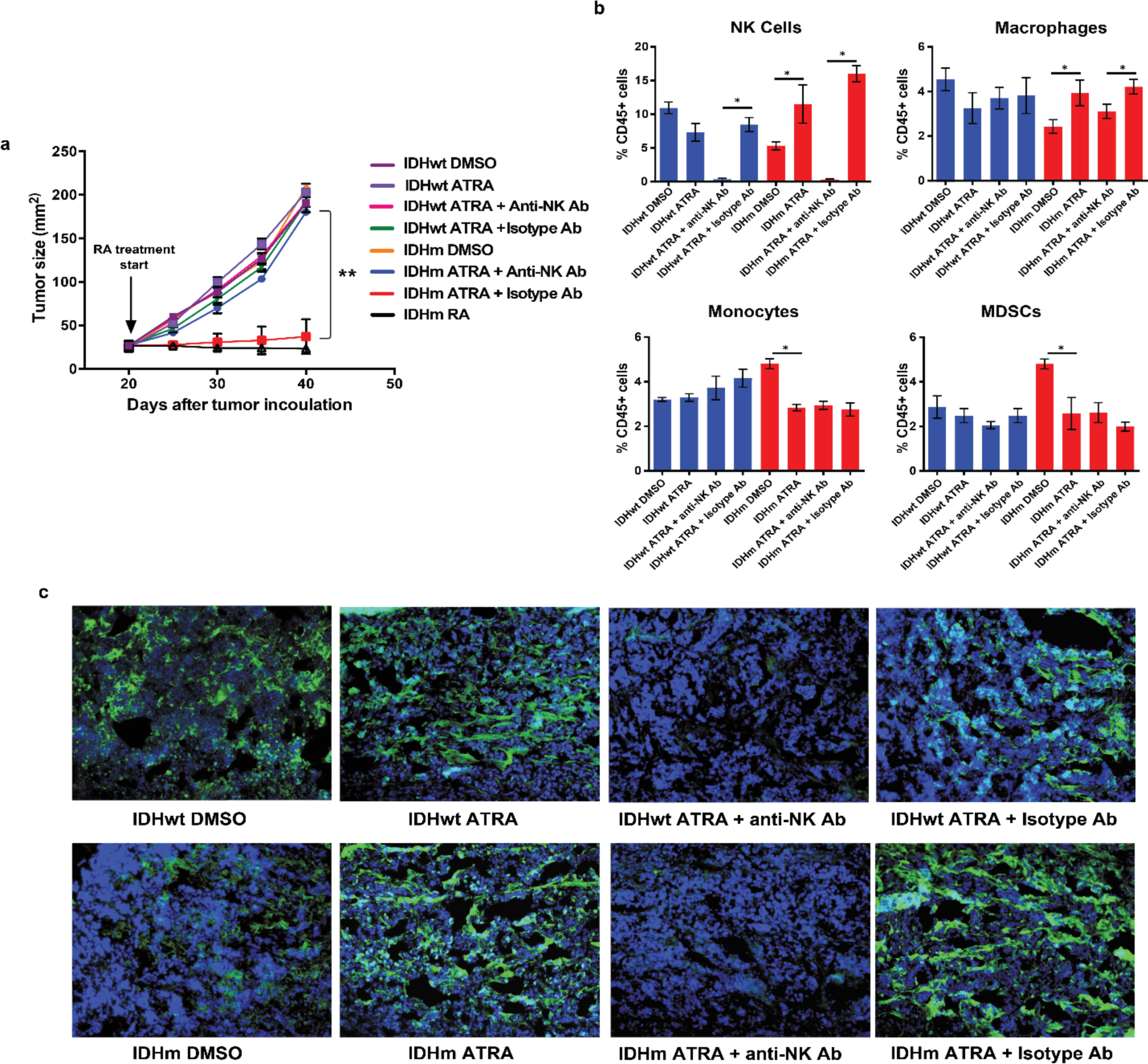
ATRA efficacy *in vivo* is dependent upon presence of NK cells. (a) BALB/c mice were implanted with IDHwt or IDHm tumor cells (as in **Extended Data Figure 5**) and treated with either 10 mg/kg ATRA or DMSO, and in some conditions 100µg of anti-NK1.1 antibody, isotype antibody, or no antibody. Tumor growth curves of mice from the different experimental groups are shown. (b) Flow cytometry analysis of immune cell infiltrates in tumor samples from all groups. (c) Immunohistochemistry of paraffin embedded sections of tumor. Nuclei are marked with DAPI (blue) while NK cells are marked with green fluorescent protein. Images shown are representative of 3 independent fields of view. Magnification 10X. (a-b) * *p* < 0.05 ** *p* < 0.001 calculated by two-tailed Student’s *t-*test.

**Extended Data Figure 7.**
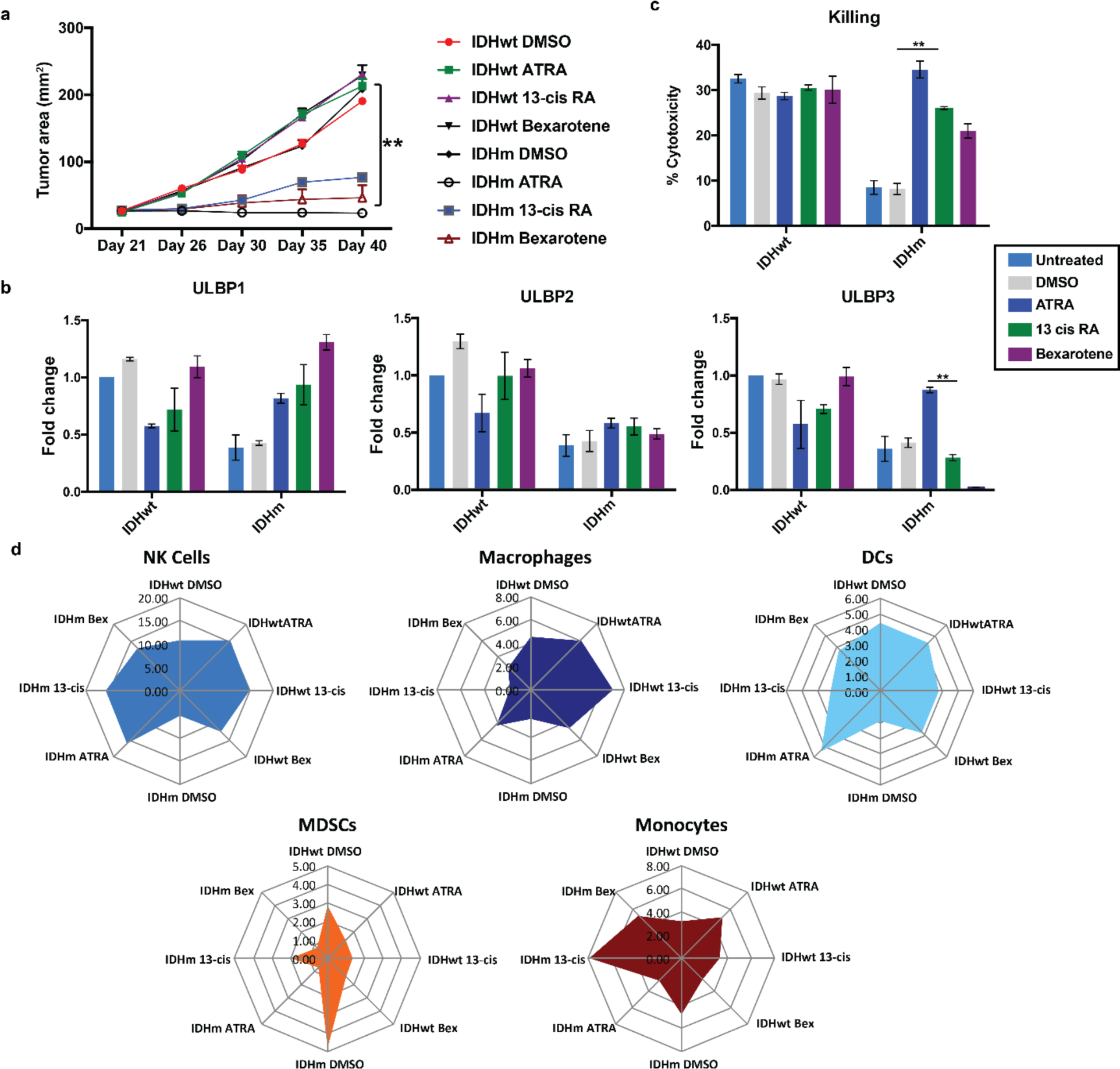
ATRA is more effective at inducing tumor stasis in IDHm tumor-bearing mice compared to other RA isoforms. (a) IDHwt or IDHm tumor-bearing mice were treated with either 10mg/kg of ATRA, 13-cis RA, or bexarotene every alternate day for 21 days. Tumor growth curves are shown. ** *p* < 0.005 calculated by two-tailed Student’s *t*-test. (b) *Ex vivo* analysis of NKG2DL gene expression in mice treated with RA isoforms or DMSO on the regimen above, assessed by qPCR. (c) Percentage of 7-AAD^+^ tumor cells after co-culture with human NK cells as measured by flow cytometry. ** *p* < 0.005 calculated by two-tailed Student’s *t*-test. (d) Flow cytometry quantification of immune cells infiltrating IDHwt or IDHm tumors that were treated with a retinoid acid derivative or DMSO.

**Extended Data Figure 8.**
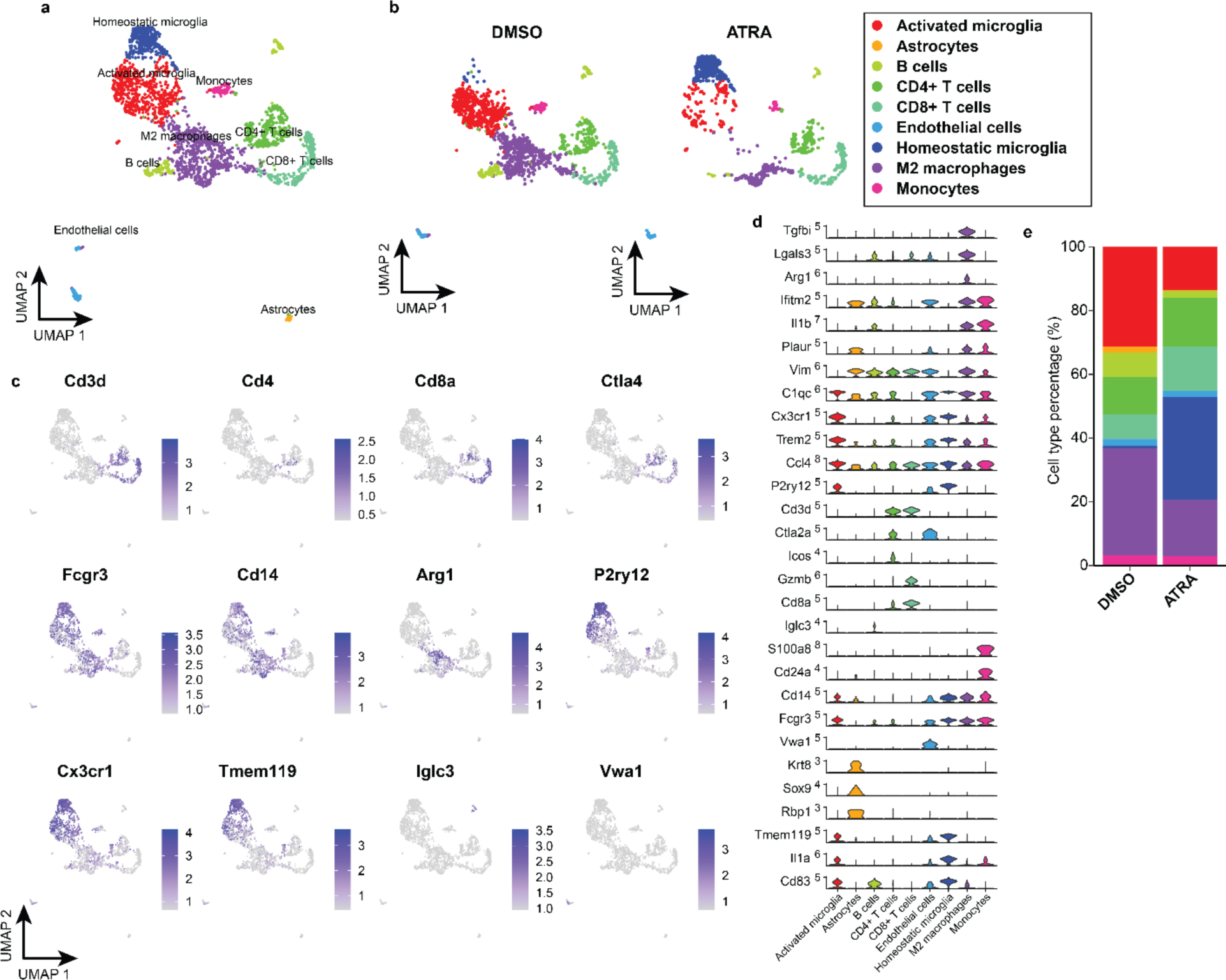
ATRA treatment alters the tumor microenvironment in mice with IDHm tumor. (a) UMAP visualization of CD45^+^ sorted intratumoral cells from ATRA-treated and DMSO-treated SB-IDHm mice (2,454 cells total, *n* = 3 pooled samples each) (b) UMAP plots of cells from (a) separated by condition, highlighting changes in the tumor immune microenvironment invoked by ATRA treatment. (c) Expression of selected marker genes overlaid on UMAP plots. (d) Violin plots of normalized expression of marker genes and genes involved in immune activation and immunosuppression in each cell population. (e) Boxplot of cell type proportions between ATRA-treated and DMSO-treated conditions.

