## Supplementary Discussion for "All-trans retinoic acid induces durable tumor immunity in IDH-mutant gliomas by rescuing transcriptional repression of the CRBP1-retinoic acid axis"

ATRA’s immunogenic potential runs well beyond altering myeloid lineages. ATRA-regulated immunomodulation maintains tissue-specific homeostasis of dendritic cells, T cells, and B cells (particularly in the gut)^1–5^. In addition, RA also upregulates the transcription of activating stress ligands that are engaged by ILC1-type cells to promote natural cytotoxicity^2^.

Consequently, complete loss of ATRA in any cell type should raise suspicion. In IDHm gliomas, loss of RA signaling is mediated by hypermethylation and transcriptional repression of *RBP1*, the product of which is a chaperone protein that transports retinol to cytoplasmic retinaldehyde dehydrogenases (particularly RDH10) for biosynthesis of ATRA^6,7^. Importantly, we found that stable expression of *RBP1* in the presence of retinol (a required ATRA precursor) reconstitutes NKG2D ligand expression and sensitizes IDHm glioma cells to NK cell-mediated cytotoxicity. Conversely, knockdown of *RBP1* expression in IDHwt glioma cells is sufficient to inhibit expression of NKG2D ligands and promote resistance to NK cells. Taken together, our data demonstrates that *RBP1* regulates NKG2D-mediated immune surveillance of glial cells by modulating intracellular RA levels and that its absence is both uniquely detrimental to anti-tumor immunity and permissive of immune evasion in IDHm gliomas.

In malignancies with profound epigenetic alterations (APML, acute myelocytic leukemia, and cutaneous T cell lymphoma) ATRA has demonstrated significant immune chemotactic effects^8^. In APML, for instance, ATRA treatment induced the migration of apoptotic tumor cells towards phagocytic cells in a CX3CL1-dependent manner^9^. It is notable that induction of anti-tumor immunity in glioma is largely specific to IDHm cells, suggesting that epigenetic dysfunction is required for ATRA-mediated immune activation. This study is the first to report the role of retinoids in mediating genotype-specific immune cell chemotaxis in solid tumors. Future studies are required to understand immune-related ATRA function and phenotypes in other IDHm solid malignancies.

Our results corroborate the impact of ATRA on the positive regulation of immune chemotaxis by defining two somewhat unexpected chemokines, CCL2 and CXCL12, as key constituents of the ATRA chemotactic machinery. CCL2 is a known transcriptional target of ATRA and modulates anti-tumor immunity or pro-tumorigenic properties in a manner dependent on tumor contexture and histology^4,10^. Paradoxically, elevated CCL2 expression levels in the tumor microenvironment of some malignancies are associated with poor prognosis^11^. In breast carcinoma, CCL2 stimulates the migration of mammary carcinoma cell lines and mediates the recruitment of suppressive monocyte populations that support the establishment of a metastatic niche^12^. Conversely, in B16 models of melanoma, tumor-induced CCL2 expression exerts vigorous anti-tumor effects by inducing chemotaxis of γδ T cells^13^. Anti-tumorigenic CCL2 effects typically correlate with increased migration of activated NK and T cells, as well as anti-tumor N1-type neutrophils^14,15^. The dichotomous role of CCL2 during immune surveillance may be a consequence of differential signaling occurring through different isoforms of its CCR2 receptor^16^, and future studies will determine the potential receptors that mediate anti-tumor immune homeostasis in IDHm glioma. Unlike CCL2, CXCL12 signaling is more consistently associated with tumor immune suppression via recruitment suppressive CD11b^+^Gr1^+^ myeloid cells, plasmacytoid dendritic cells, and intratumoral Tregs - all of which impede adaptive anti-tumor immune responses^17–19^. Thus, ATRA-mediated repression of CXCL12 production likely contributes to the pro-inflammatory and anti-tumor immune responses seen in IDHm tumors.
