## Supplementary Tables 4-7 for "All-trans retinoic acid induces durable tumor immunity in IDH-mutant gliomas by rescuing transcriptional repression of the CRBP1-retinoic acid axis"

**Supplemental Table 1: Differentially expressed genes after ATRA treatment in IDHm astrocytoma cell lines.**

**Supplemental Table 2: Differentially expressed genes after ATRA treatment in IDHm astrocytoma cell lines.**

**Supplemental Table 3: Differentially expressed genes after ATRA treatment in IDHm astrocytoma cell lines.**

**Supplemental Table 4: Cell line information**

| Specimen | IDH status | WHO Grade | 1p LOH | 19q LOH | 10q LOH | 17p LOH | P53 mutation | EGFR mutation | EGFR amplification | IDH2 |
| --- | --- | --- | --- | --- | --- | --- | --- | --- | --- | --- |
| UP612 | IDH1- R132H | 2 | + | + | - | - | - | - | - | - |
| UP416 | IDH1- R132H | 3 | + | + | - | - | - | - | - | - |
| UP807 | IDH1- R132H | 2 |  |  |  |  |  |  |  |  |
| UP205 | IDH1 R132H | 3 | + | + | - | - | - | - | - | - |
| TS603 | IDH1 R132H | 3 |  |  |  |  |  |  |  |  |
| UP319 | WT | 4 | - | - | + | - | + | - | - | - |
| UP226 | WT | 4 | - | - | + | + | - | - | + | - |
| UP607 | WT | 4 | - | - | - | - | - | - | - | - |
| TS667 | WT | 3 |  |  |  |  |  |  |  |  |
| UP101 | WT | 4 |  |  |  |  |  |  |  |  |

Annotation of IDH1 mutation status and WHO grade for cells lines used in this study. Additional histopathological testing included where available.

**Supplemental Table 5: Patient Demographics**

| AGE | WHO Grade | Retinoids | Concurrent Therapy | PFS |
| --- | --- | --- | --- | --- |
| 48 | III | 13-Cis | CCNU | 65 |
| 46 | III | 13-Cis | PCV | 55 |
| 43 | II | 13-Cis | PCV | 93 |
| 36 | II | 13-Cis | None | 94 |
| 45 | III | ATRA | None | 32 |
| 45 | III | ATRA | Avastin | 851 |
| 69 | III | ATRA | None | 167 |
| 57 | II | ATRA | None | 206 |
| 43 | II | ATRA | None | 991 |

Eight patients with multiple recurrent WHO grade II, III gliomas were treated with either 13-Cis or ATRA based on prescriber preference in a non-blinded, non-randomized fashion. One patient was initially treated with 13-Cis and subsequently had progression of disease and was later treated with ATRA. Progression free survival (PFS) was determined by evidence of progression on MRI.

**Supplementary Table 6: Flow Cytometry Gating Strategy**

**Gating Strategy**

| **Cell type** | **Gating strategy** |
| --- | --- |
| All immune cells | CD45+ |
| All T cells | CD45+ CD3+ |
| CD4+ T cells | CD45+ CD3+ CD4+ |
| Tregs | CD45+ CD3+ CD4+CD25+ FoxP3+ |
| CD8+ T cells | CD45+ CD3+ CD8+ |
| Activated CD8+ cells | CD45+ CD3+ CD8+IFNg+ |
| NK cells | CD45+ CD3- Nkp46+ CD49b+ |
| NKT cells | CD45+ CD3- Nkp46+ CD49b+CD8+ |
| B cells | CD45+ CD19+ B220+ |
| All Macrophages | CD45+ CD11b+ F4/80+ |
| M1 Macrophages | CD45+ CD11b+ F4/80+ CD80+ CD86+ MHC-II + |
| M2 Macrophages | CD45+ CD11b+ F4/80+ CD206+ VISTA+ |
| DCs | CD45+ CD11c+ MHC-II+ |
| Monocytes | CD45+ CD11b+ Gr1- CD115+ |
| Mono-MDSCs | CD45+ CD11b+ Gr1+ Ly6C + |
| PMN-MDSCs | CD45+CD11b+ Gr1+ Ly6G+ |

**Supplementary Table 7: Flow Cytometry Antibody Panels**

**NK/B/T-cell Panel:**

| **Marker** | **Fluorochrome** | **Brand** | **Cat #** |
| --- | --- | --- | --- |
| CD45 | PE-Cy7 | Invitrogen | 25-0451-81 |
| CD3 | APC-Cy7 | Biolegend | 100330 |
| CD4 | PerCP-Cy5.5 | Invitrogen | 45-0042-80 |
| CD8 | SB645/BV650 | Invitrogen | 64-0081-80 |
| NKp46 | FITC | BdBioscience | 560756 |
| DX5 (CD49b) | APC | Invitrogen | 17-5971-81 |
| B220 | PE | Invitrogen | 11-3351-80 |
| CD25 | BV421/eFluor450 | BdPharmigen | 562606 |
| FoxP3 | PE-eFluuor610 | Invitrogen | 61-5773-82 |
| CD19 | AlexaFluor700 | Invitrogen | 56-0193 |
| IFNg | BV711 | BdHorizon | 564336 |

**Macrophage Panel:**

| **Marker** | **Fluorochrome** | **Brand** | **Cat #** |
| --- | --- | --- | --- |
| CD45 | PE-Cy7 | Invitrogen | 25-0451-81 |
| CD11b | APC | Invitrogen | 17-0112-81 |
| CD80 | PerCP-Cy5.5 | Biolegend | 104722 |
| CD86 | BV786 | BD | 740877 |
| F4/80 | eFluor450 | Invitrogen | 48-4801-80 |
| MHC-II (I-Ab/d) | FITC | Invitrogen | 11-5321-81 |
| CD206 | PE | Invitrogen | 12-2061-82 |
| VISTA | BV711 | BDOptiBuild | 742724 |

**Monocyte/MDSC Panel:**

| **Marker** | **Fluorochrome** | **Brand** | **Cat #** |
| --- | --- | --- | --- |
| CD45 | PE-Cy7 | Invitrogen | 25-0451-81 |
| CD11b | APC | Invitrogen | 17-0112-81 |
| CD11c | PerCP-Cy5.5 | Invitrogen | 45-0114-80 |
| Gr1 | AlexaFluor700 | Invitrogen | 56-5931-82 |
| LyC | eFluor780(APC-Cy7) | Invitrogen | 47-5932-80 |
| LyG | PE-eFluor610 | Invitrogen | 619668-80 |
| CD115 | PE | Invitrogen | 12-1152-81 |
| MHC-II (I-Ab/d) | FITC | Invitrogen | 11-5321-81 |
